# Dextrin conjugation to colistin inhibits its toxicity, cellular uptake and acute kidney injury *in vivo*

**DOI:** 10.1101/2023.11.02.565265

**Authors:** Mathieu Varache, Siân Rizzo, Edward J. Sayers, Lucy Newbury, Anna Mason, Chia-Te Liao, Emilie Chiron, Nathan Bourdiec, Adam Jones, Donald J. Fraser, Philip R. Taylor, Arwyn T. Jones, David W. Thomas, Elaine L. Ferguson

## Abstract

The acute kidney injury (AKI) and dose-limiting nephrotoxicity, which occurs in 20-60% of patients following systemic administration of colistin, represents a challenge in the effective treatment of multi-drug resistant gram-negative infections. To reduce clinical toxicity of colistin and improve targeting to infected /inflamed tissues, we previously developed dextrin-colistin conjugates, whereby colistin is designed to be released by amylase-triggered degradation of dextrin in infected and inflamed tissues, after passive targeting by the enhanced permeability and retention effect. Whilst it was evident *in vitro* that polymer conjugation can reduce toxicity and prolong plasma half-life, without significant reduction in antimicrobial activity of colistin, it was unclear how dextrin conjugation would alter cellular uptake and localisation of colistin in renal tubular cells *in vivo*. We discovered that dextrin conjugation effectively reduced colistin’s toxicity towards human kidney proximal tubular epithelial cells (HK-2) *in vitro*, which was mirrored by significantly less cellular uptake of Oregon Green (OG)-labelled dextrin-colistin conjugate, when compared to colistin. Using live-cell confocal imaging, we revealed localisation of both, free and dextrin-bound colistin in endolysosome compartments of HK-2 and NRK-52E cells. Using a murine AKI model, we demonstrated dextrin-colistin conjugation dramatically diminishes both proximal tubular injury and renal accumulation of colistin. These findings reveal new insight into the mechanism by which dextrin conjugation can overcome colistin’s renal toxicity and show the potential of polymer conjugation to improve the side effect profile of nephrotoxic drugs.

**Graphical abstract:** 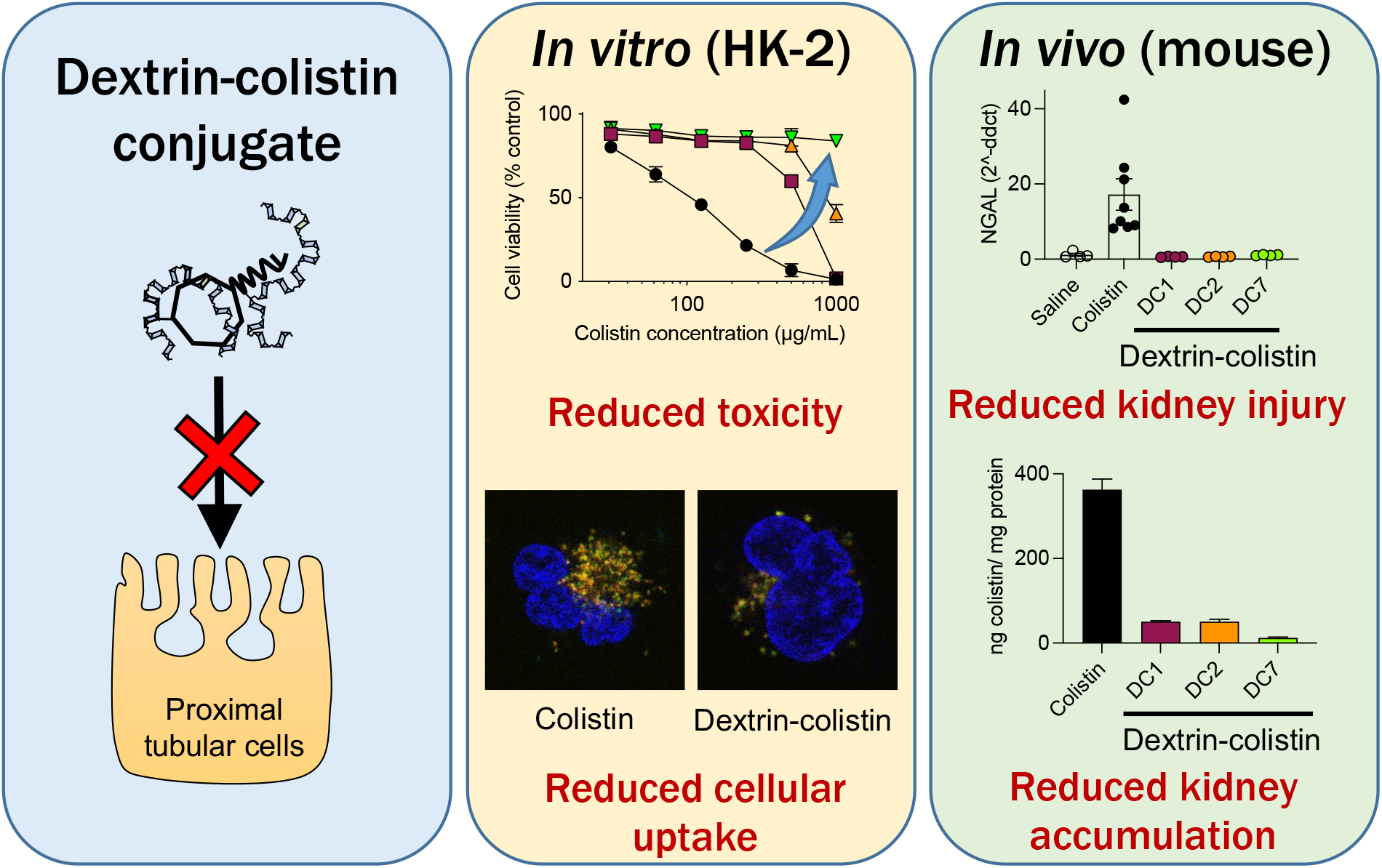

## Introduction

Polymyxins are highly effective antibiotics in the treatment of multi-drug resistant (MDR) infections caused by gram-negative bacteria. The reported incidence of nephrotoxicity in patients receiving these drugs (26.7% for colistin and 29.8% for polymyxin B, according to a recent meta-analysis)^1^ has led researchers to develop alternative dosing strategies and delivery systems to reduce this toxicity without compromising antimicrobial activity.

Until recently, the exact mechanism of polymyxin nephrotoxicity and its distribution in the kidneys was poorly understood. Recent research by Yun et al. has demonstrated that polymyxin B distributes primarily to the proximal tubular cells of the renal cortex,^2^ due to extensive reabsorption and accumulation, where it accumulates at concentrations of up to 4,760-fold higher than the extracellular compartment.^3^ Internalisation of colistin at the luminal side of renal proximal tubule cells is driven by endocytosis (principally via the megalin receptor)^4^ and, due to its polycationic nature, facilitative transport (via the human peptide transporter 2 (PEPT2) and carnitine/organic cation transporter 2 (OCTN2)).^5, 6^ Following internalisation, colistin induces dysfunction of the mitochondria and endoplasmic reticulum (ER), inducing concentration- and time-dependent cellular apoptosis by activation of the death receptor, mitochondrial and ER pathways.^7^ Histopathological damage, characterised by tubular dilation, epithelial cell vacuolisation and necrosis (in the absence of inflammatory response or interstitial fibrosis)^2, 3, 7, 8^ is also observed. More recently, a direct interaction between colistin and the phospholipid bilayer of renal tubular cells has been proposed to contribute to polymyxin-induced nephrotoxicity.^9^ Even intravenous (IV) administration of the pro-drug, colistimethate sodium (CMS), has been observed to induce cumulative dose- and duration-dependent increases in serum creatinine (a marker of kidney damage).^10, 11^

We previously described the potential use of dextrin-colistin conjugation, designed to improve targeting to sites of infection by amylase-triggered degradation of the polymeric carrier (using polymer-masked unmasked protein therapy (PUMPT), as an attempt to reduce the observed nephrotoxicity.^12–15^ These studies demonstrated that dextrin-colistin conjugates had comparable antibacterial activity to Colomycin^®^, but with reduced *in vitro* toxicity and prolonged plasma half-life in rats (after a single dose). Hypothesising that dextrin conjugation would also potentially reduce renal endocytosis, accumulation and ultimately, nephrotoxicity, we investigated the *in vitro* and *in vivo* nephrotoxicity of dextrin-colistin conjugates. The internalisation and cellular localisation of Oregon Green (OG) 488-labelled conjugates was assessed in HK-2 and NRK-52E renal proximal tubule cells using flow cytometry and confocal microscopy, and *in vitro* cytotoxicity was assessed by measuring necrosis, metabolic activity and caspase activity. The dextrin-colistin conjugates were studied in a murine model of colistin-induced AKI to characterise the renal changes induced at the cellular and renal function level. Remarkably, we established that dextrin conjugation effectively prevents colistin-induced AKI, and demonstrated reduced cellular uptake of the antibiotic by proximal tubular cells. This work highlights, for the first time, the potential of polymer conjugation as a valuable tool to improve the side effect profile of nephrotoxic drugs.

## Materials and Methods

### Materials

Type I dextrin from corn (M_w_ = 7,500 g/mol, degree of polymerisation = 50), colistin sulfate, N-hydroxysulfosuccinimide (sulfo-NHS) and dimethyl sulfoxide (DMSO) were purchased from Sigma-Aldrich (Poole, UK). QIAzol reagent and miRNeasy kit were from Qiagen (Manchester, UK). BCA protein assay kit, N,N′-dicyclohexyl carbodiimide (DCC), 1-ethyl-3- (3-(dimethylamino)propyl carbodiimide hydrochloride) (EDC), Gibco^TM^-branded keratinocyte serum-free medium (K-SFM) with L-glutamine, epidermal growth factor (EGF), bovine pituitary extract (BPE), Dulbecco’s modified Eagle’s medium (DMEM) with high glucose (4.5 g/L) and GlutaMAX^TM^, fetal bovine serum (FBS), 0.05% w/v trypsin-0.53 mM EDTA, Oregon Green^TM^ 488 (OG) cadaverine, OG carboxylic acid, wheat germ agglutinin-Alexa 594 (WGA-AF594), leupeptin, Hoechst 33342, BODIPY^TM^ TR Ceramide complexed to bovine serum albumin (BSA), Mitotracker^TM^ Red FM, high capacity cDNA kit and power SYBR green were obtained from ThermoFisher Scientific (Loughborough, UK). Narrow (polyethylene oxide, Mw = 24,063 g/mol, PDI = 1.02) and broad (dextran, Mw = 68,162 g/mol, PDI = 1.24) PolyCAL^TM^ standards were purchased from Malvern Panalytical (Malvern, UK). Unless otherwise stated, all chemicals were of analytical grade and used as supplied. All solvents were of general reagent grade (unless stated) and were from Fisher Scientific (Loughborough, UK).

### Cell Culture

Human kidney proximal tubule cells (HK-2) were from ATCC (Manassas, USA) and cultured in K-SFM. Rat proximal tubule cells (NRK-52E) were from ECACC (Salisbury, UK) and cultured in DMEM with 5% v/v FBS. Cells were screened to be free of mycoplasma contamination upon thawing and monthly thereafter. Cell lines were maintained in log-phase proliferation at 37°C with 5% CO_2_ in their respective culture medium and passaged using 0.05% w/v trypsin-0.53 mM EDTA.

### Animals

All animal experiments were conducted according to the United Kingdom Use of Animals (Scientific Procedures) Act 1986. Animal work was reviewed by the Animal Welfare and Ethical Review Body under the Establishment Licence held by Cardiff University and authorised by the UK Home Office. C57BL/6 mice (male, 8 weeks) were purchased from Charles River Laboratories (Bristol, UK). The mice were given a 7-day period of acclimatisation to their new surroundings and were housed and handled according to the local institutional policies.

### Synthesis and characterisation of dextrin-colistin conjugates and OG-labelled probes

Dextrin-colistin conjugates, containing dextrin with 1, 2.5 and 7.5 mol% succinoylation, were synthesised using EDC and sulfo-NHS and characterised as previously described.^15^ Free colistin content, analysed by fast protein liquid chromatography (FPLC), was determined to be <3%. The characteristics of dextrin-colistin conjugates used in these studies are summarised in Table 1.

**Table 1.** Characteristics of dextrin-colistin conjugates used in these studies.

| Conjugate name | Dextrin succinylation (mol%) | Mw<sup>a</sup> (Mw/Mn) | R<sub>H</sub><sup>a</sup> (nm) | Protein content (% w/w) | Molar ratio (x dextrin: 1 colistin) | Free protein (%) |
| --- | --- | --- | --- | --- | --- | --- |
| DC1 | 1 | 29,600 (1.6) | 2.6 | 5.8 | 2.9 | 2.1 |
| DC2 | 2.5 | 30,300 (1.5) | 2.7 | 6.4 | 2.6 | 2.4 |
| DC7 | 7.5 | 59,800 (1.8) | 3.2 | 14.0 | 1.1 | 0.7 |

<sup>a</sup> measured using Viscotek TDA 302 detector with 2 × Ultrahydrogel 250 columns. Mw Molecular weight, Mn Molecular number, R<sub>H</sub> Radius of hydration

To enable visualisation of conjugates by flow cytometry and confocal microscopy, succinoylated dextrin, colistin and dextrin-colistin conjugate were fluorescently labelled with Oregon Green 488 (OG) and characterised according to previously published methods,^16^ including spectrophotometric and fluorometric analysis (see Supplementary data for detailed description).

### Evaluation of *in vitro* cytotoxicity

Cytotoxicity of colistin sulfate and dextrin-colistin conjugates was assessed in HK-2 and NRK-52E cell lines using a multiplexed system that measures cell viability (CellTiter-Blue® (CTB) cell viability assay kit), membrane integrity (CytoTox-One^TM^ homogeneous membrane integrity assay kit: lactate dehydrogenase (LDH release) and caspase activity/ apoptosis (Caspase-Glo® 3/7 assay system kit) (all from Promega, WI, USA), as described previously^17^ (see Supplementary data for detailed description).

### Determination of uptake by flow cytometry

To study cellular uptake, cells were first seeded in 6-well plates (500,000 cells/mL) in 1 mL CM and allowed to adhere for 24 h at 37°C with 5% CO_2_. Cell association experiments at 37°C were conducted under normal cell culture conditions, but for experiments at 4°C, cells were pre-incubated on ice for 30 min prior to addition of the probe. Solutions of dextrin-OG, colistin-OG or dextrin−colistin-OG were freshly prepared at sub-toxic colistin concentrations in CM (1.5 μg OG base/mL), filter-sterilised (0.22 μm), then equilibrated to either 37 or 4°C for 30 min. Cell culture medium was removed from each well and replaced with probe solutions (1 mL/ well), before incubating the microplates for 1 h at 4 or 37°C. Subsequently, cell culture medium was removed from each well and analysed for free OG content by PD-10 chromatography, as previously described.^18^ To analyse cellular uptake, the original microplates were placed on ice before washing with ice-cold PBS (3 x 1 mL). After the final wash, trypsin-EDTA (200 μL) was added to each well and the cells harvested into falcon tubes. Each tube was topped up to 1 mL with PBS then centrifuged twice at 4°C (250 x *g* for 5 min). Finally, cells were resuspended in ice-cold PBS (200 μL) and analysed using a Becton Dickinson FACS Canto II cytometer equipped with an argon laser (488 nm) and emission filter for 550 nm. Data were collected for 10,000 cell counts per sample and processed using BD FACSDiva^TM^ software version 6.1.3. Control cells incubated with medium only were used to establish background fluorescence and define fluorescence gated regions. Results were corrected for cell autofluorescence and expressed as (geometric mean × % positive cells)/100, where positive cells are those cells falling within the region of positive staining (Figure 2a,b). Internalisation was calculated by subtracting the cell-associated fluorescence at 4°C (extracellular binding) from that at 37°C (intracellular uptake plus extracellular binding), and % internalisation was derived from the internalised OG-labelled probes compared to the total cell-associated fluorescence (at 37°C). Cells were plated in triplicate (n=3 technical replicates) and each experiment was repeated 3 times (n=3 biological replicates). Internalisation was expressed as mean ± SEM.

### Determination of intracellular fate

To study endocytic trafficking, HK-2 and NRK-52E cells were, respectively, seeded into tissue culture-treated 35 mm plastic dishes from Ibidi (Glasgow, UK) or uncoated 35 mm glass-bottom dishes (No. 1.5 coverslip, 10 mm glass diameter) from MatTek (Ashland, USA). HK-2 and NRK-52E cells were seeded in 1.5 mL of CM at 60,000 and 75,000 cells/dish, respectively. After 24 h incubation at 37°C with 5% CO_2_, cell culture medium was removed from each dish and replaced with CM containing wheatgerm agglutinin-Alexa Fluor 594 (WGA-AF594; 0.1 or 5 µg/mL for HK-2 and NRK-52E cells, respectively), as a physiological marker for endolysosomal content.^16^ After 4h, cell culture medium was removed and cells washed with PBS (3 x 1.5 mL). After the final wash, for the short chase experiment, cell culture medium was removed and replaced with freshly prepared filter-sterilised (0.22 μm) CM containing dextrin-OG, colistin-OG or dextrin-colistin-OG (5-10 μg/mL OG base) and leupeptin (200 µM). After a further 2 h incubation (pulse), cells were intensively washed with PBS, then, for the short chase experiment, cells were immediately incubated with Hoechst 33342 solution (1 µg/mL) in CM (without phenol red or leupeptin) for 20 min before imaging. In contrast, for the long chase experiment, after incubation with WGA-AF594 for 4 h, cells were washed and 1.5 mL CM containing leupeptin (200 µM) was added to each well. After 1 h incubation, media was removed and replaced with CM containing filter-sterilised (0.22 μm) dextran-Alexa Fluor 488 (dextran-AF488, as control), dextrin-OG, colistin-OG or dextrin-colistin-OG and leupeptin (200 µM) for 2 h. Subsequently, cells were intensively washed with PBS then CM containing leupeptin (200 µM) was added and the plates incubated at 37°C with 5% CO_2_ overnight. Finally, cells were washed with PBS then incubated with Hoechst 33342 solution (1 µg/mL) in CM (without phenol red or leupeptin) for 20 min before imaging.

To study accumulation in mitochondria and Golgi in HK-2 cells, the same method was used, except that the WGA-AF594 step was omitted and at the end of the chase period, cells were incubated with Mitotracker^TM^ Red FM (100 nM) or BODIPY^TM^ TR ceramide (5 µM) in CM for 20 min before washing with PBS (3 x 1.5 mL). The protocols used here are summarised in Scheme S1,2.

### Live-cell imaging by confocal microscopy

Confocal microscopy of cells treated as above was performed using a Leica SP5 laser scanning system (37°C with 5% CO_2_). Confocal imaging was performed sequentially with the 405 nm, 488 nm, 543 nm and 633 nm lasers and using a 1.4 N/A 63x oil immersion CSII objective at 1,000 Hz with a line average of 3 (bi-directional scanning) and the pinhole set to 1 airy unit. Sequential scans were performed to minimise bleed from one channel to another and sample photobleaching. Images were acquired with a raster size of 1,024 x 1,024 and a zoom of 2.5 to give an apparent voxel size of 130 x 130 x 500 nm (XYZ); here the pixel size is smaller than the resolution limit attainable by the microscope. At least, eight representative images (single section) were obtained from each sample; representative images for each condition are shown. Images were analysed and processed using ImageJ^19^ and the JACoP^20^ plugin for ImageJ, using Pearson’s coefficient to determine co-localisation of OG-labelled probes with WGA-AF594. Thresholds for images were automatically determined using the Otsu thresholding algorithm, and the co-localisation value returned was used to calculate the mean co-localisation of 8 images.

### *In vivo* acute kidney injury model

To confirm an appropriate dose of colistin sulfate that could achieve significant acute kidney injury without general toxicity, phase 1 involved random division of mice into two groups (n = 4 animals per group), as follows: control (saline), colistin sulfate. In phase 2, mice were randomly divided into four groups (n = 4 animals per group), as follows: colistin sulfate, DC1, DC2 and DC7 (40 mg colistin base/kg/day). To avoid experimental bias in subsequent analyses, each group of mice was fitted with an ear tag (right only, left only, right & left or none) and assigned an identification code that was only decrypted after sample analysis was complete. Colistin sulfate and dextrin-colistin conjugates were administered intraperitoneally every 12 h for 7 consecutive days (280 mg colistin equiv./kg total dose). An equivalent volume of saline was used as control. 12 h after the last dose, mice were euthanised by schedule 1 using CO_2_, followed by cardiac puncture. Blood samples were collected, and the serum was separated by centrifugation (2000 *g* for 10 min) and stored at −80°C until assayed. Following perfusion with ice-cold PBS (20 mL) via the left ventricle, the kidneys and tissue samples were harvested and snap-frozen for subsequent biochemical, ELISA and quantitative reverse transcription–PCR (RT–qPCR) studies or fixed in formalin and embedded in paraffin prior to histopathological analysis.

### Quantification of creatinine and urea

Serum creatinine and urea were determined using Alinity creatinine and urea nitrogen reagent kits (Abbott Laboratories, Abbott Park, USA), respectively, according to the manufacturer’s instructions.

### RNA extraction, RT-qPCR of KIM-1 and NGAL from mouse kidneys

Kidney samples (one quarter) were homogenised in QIAzol reagent (1 mL) in a 2 mL safe lock tube with stainless steel beads using a TissueLyser II homogeniser (Qiagen, UK) at 30 Hz until fully homogenised (1-2 min). Kidney homogenates were diluted with QIAzol (1 mL) and aliquoted into a 1.5 mL tube. Chloroform (200 µL) was added to each sample, tubes were inverted and the samples were centrifuged at 12,000 *g* for 15 min at 4°C. Each aqueous phase was then transferred to a fresh 1.5 mL tube and samples processed as per instructions for miRNAeasy kits for total RNA extraction. RNA was quantified using a nano drop, samples were diluted and 1000 ng was added to mRNA RT reactions. RT reactions were carried out using the high-capacity cDNA kit as recommended by the manufacturer. For NGAL and KIM-1, primers (Table S1) were designed using Primer-BLAST and qPCR performed using power SYBR green according to the manufacturer’s protocol. Data were analysed using the 2–ΔΔCt method.

### Histopathological examination of renal damage

Kidney tissue samples were fixed in 10% formalin, embedded in paraffin, then sections were stained by hematoxylin and eosin (H&E). Scoring of histological damage in anonymised sections was performed under light microscopy using the EGTI (Endothelial, Glomerular, Tubular, Interstitial) scoring system.^21^

### Measurement of colistin tissue levels

Kidney, liver and brain samples were homogenised in PBS (1-3 mL, adjusted to tissue weight) using gentleMACS™ M tubes (Miltenyi Biotec Ltd, Bisley, Surrey) and gentleMACS™ octo dissociator using the standard protein program. Total protein content of the homogenates was determined by BCA assay using a BSA standard calibration curve. Total tissue colistin concentration was assessed using a commercial Europroxima colistin competitive enzyme immunoassay test kit (R-Biopharm Nederland B.V, Arnhem, NL) according to the manufacturer’s instructions. Samples (n = 4) for each treatment were each evaluated in duplicate, corrected for no-cell background, then expressed as mean ng colistin/ mg of protein ± SEM.

### Statistical Analysis

Data are expressed as mean ± error, calculated as either standard deviation (SD) where *n* = 3 or standard error of the mean (SEM) where *n* > 3. Statistical significance was indicated by *, where * *p* < 0.05, ** *p* < 0.01, *** *p* < 0.001 and **** *p* < 0.0001. Evaluation of significance was achieved using a one-way analysis of variance (ANOVA) followed by Bonferroni *posthoc* tests that correct for multiple comparisons. All statistical calculations were performed using GraphPad Prism 9.2.0 for MacOS, 2021.

## Results

### Evaluation of *in vitro* cytotoxicity

Colistin sulfate caused a dose-dependent decrease in HK-2 cell viability that was greater than any of the dextrin-colistin conjugates tested (Figure 1). In contrast, the free antibiotic had almost no effect on NRK-52E cells, even at the highest concentration tested. In both cell lines, increasing the degree of dextrin succinoylation reduced the conjugates’ cytotoxicity (DC7 > DC2 > DC1) (Figure 1). Under the assay conditions, DC7 was not toxic towards HK-2 cells, whereas a 20-40% reduction in NRK-52E cell viability was seen at the highest concentration of DC7 tested (1000 µg/mL colistin base). Prolonged incubation of cells with the treatments caused more pronounced effects on cell viability.

**Figure 1.**
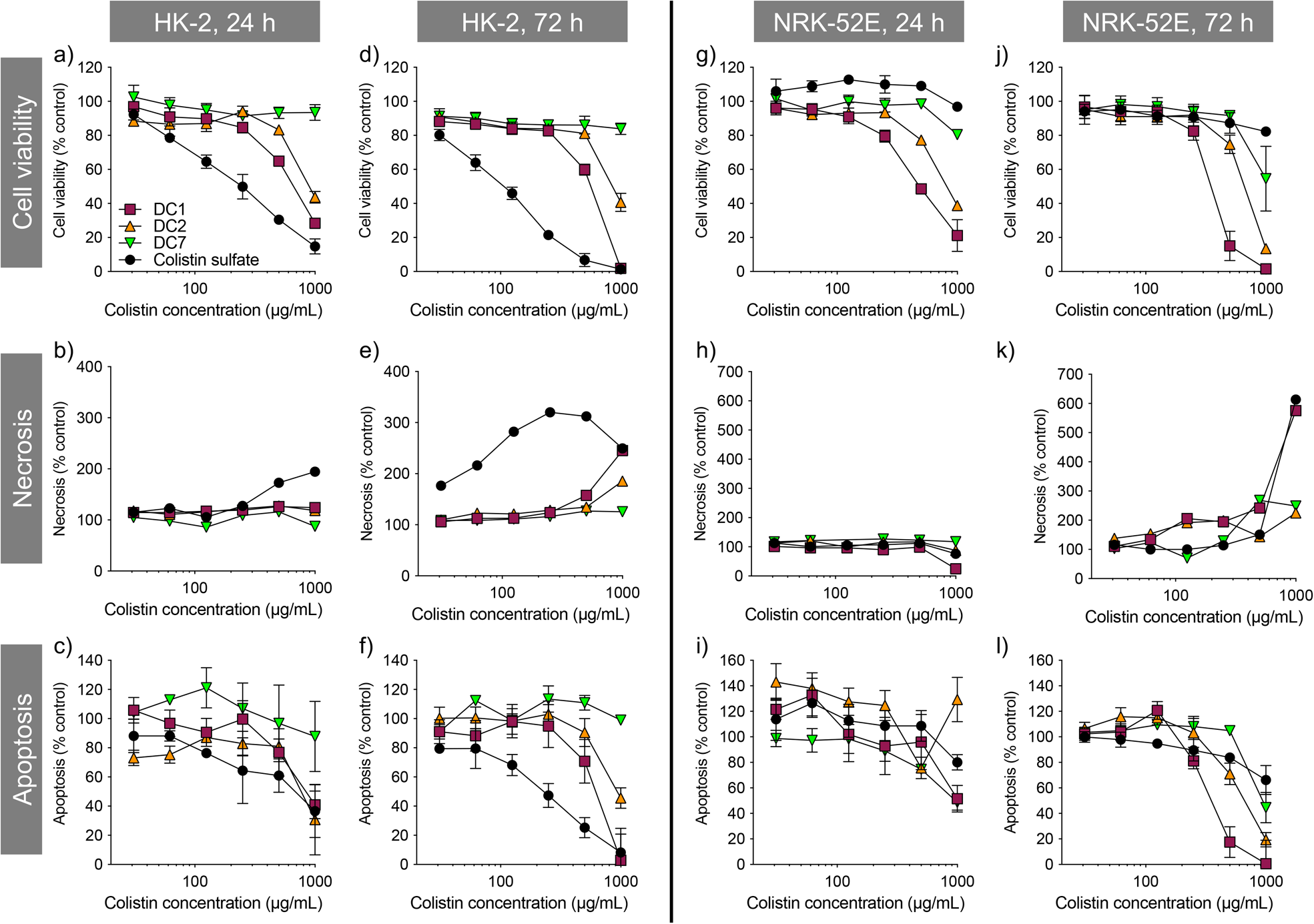
Metabolic activity (CellTiter-Blue®), necrosis (LDH release) and apoptosis (caspase 3/7 activity) of (a-f) HK-2 and (g-l) NRK-52E cells incubated for 24 or 72 h with colistin sulfate and dextrin-colistin conjugates, as a percentage of untreated (media only) controls. Data represent mean ± SD (*n*=3). Where error bars are invisible, they are within the size of data points. *μg/mL* milligram per millilitre

Colistin sulfate induced some concentration- and time-dependent loss of cell membrane integrity in both cell lines. DC1 caused necrosis in both cell lines after 72 h incubation at the highest treatment concentration, which was more pronounced than either, DC2 or DC7. No evidence of caspase 3/7-mediated apoptosis was observed for any of the treatments tested in either cell line; caspase 3/7 activity was proportional to cell viability.

### Evaluation of cellular uptake and intracellular fate

OG-labelled colistin, dextrin and dextrin-colistin conjugates were synthesised, containing <2% free OG. The characteristics of all OG-labelled probes used in these studies are summarised in Table 2 and UPLC analysis of OG-labelled colistin is shown in Figs. S1 and S2 and Table S2.

**Table 2.** Characteristics of OG-labelled probes used in these studies.

| Compound | Molar ratio (colistin or dextrin:OG) | Labelling efficiency (µg OG/mg compound)<sup>a</sup> |
| --- | --- | --- |
| Colistin | 1:0.09 | 0.76 |
| Dextrin | 1:0.14 | 8.62 |
| Dextrin-colistin | 1:0.16 (dextrin:OG)<br>1:0.17 (colistin:OG) | 8.84 |

<sup>a</sup> measured by analysis of SEC fractions (PD-10 column) for fluorescence. OG Oregon green

Cellular uptake of OG-labelled colistin was observed in both, human and rat kidney cell lines, with minimal cell-associated fluorescence at 4°C (external binding) (Figure 2). The internalisation of fluorescent probes was higher in HK-2 cells than NRK-52E cells, but both cell lines showed similar patterns of calculated internalisation (colistin > dextrin = dextrin-colistin conjugate). Analysis by SEC of the cell culture medium after incubation of cells with fluorescent probes did not show any detectable free OG after incubation for 4 h (Figure 2e,f). Live-cell imaging of HK-2 cells incubated with fluorescently-labelled compounds showed that dextran-AF488 (as a positive control for lysosomal accumulation), OG-labelled dextrin, dextrin-colistin conjugate and colistin fluorescence was almost exclusively localised in WGA-AF594-positive vesicles (Figure 3, S3). Labelling either the Golgi, via BODIPY-ceramide, or the mitochondria, via MitoTracker, in live HK-2 cells showed a marked separation of fluorescence localisation between these organelles and any of the OG-labelled compounds (Figure S4, S5). In contrast, live-cell imaging of NRK-52E cells incubated with fluorescently-labelled compounds showed negligible co-localisation of OG-labelled compounds with WGA-AF594 (Figure S6, S7). Dextrin conjugation significantly reduced co-localisation of OG-labelled colistin with the endolysosomal marker, WGA-AF594 (short chase, *p*<0.01; long chase, *p*<0.0001) (Figure S7).

**Figure 2.**
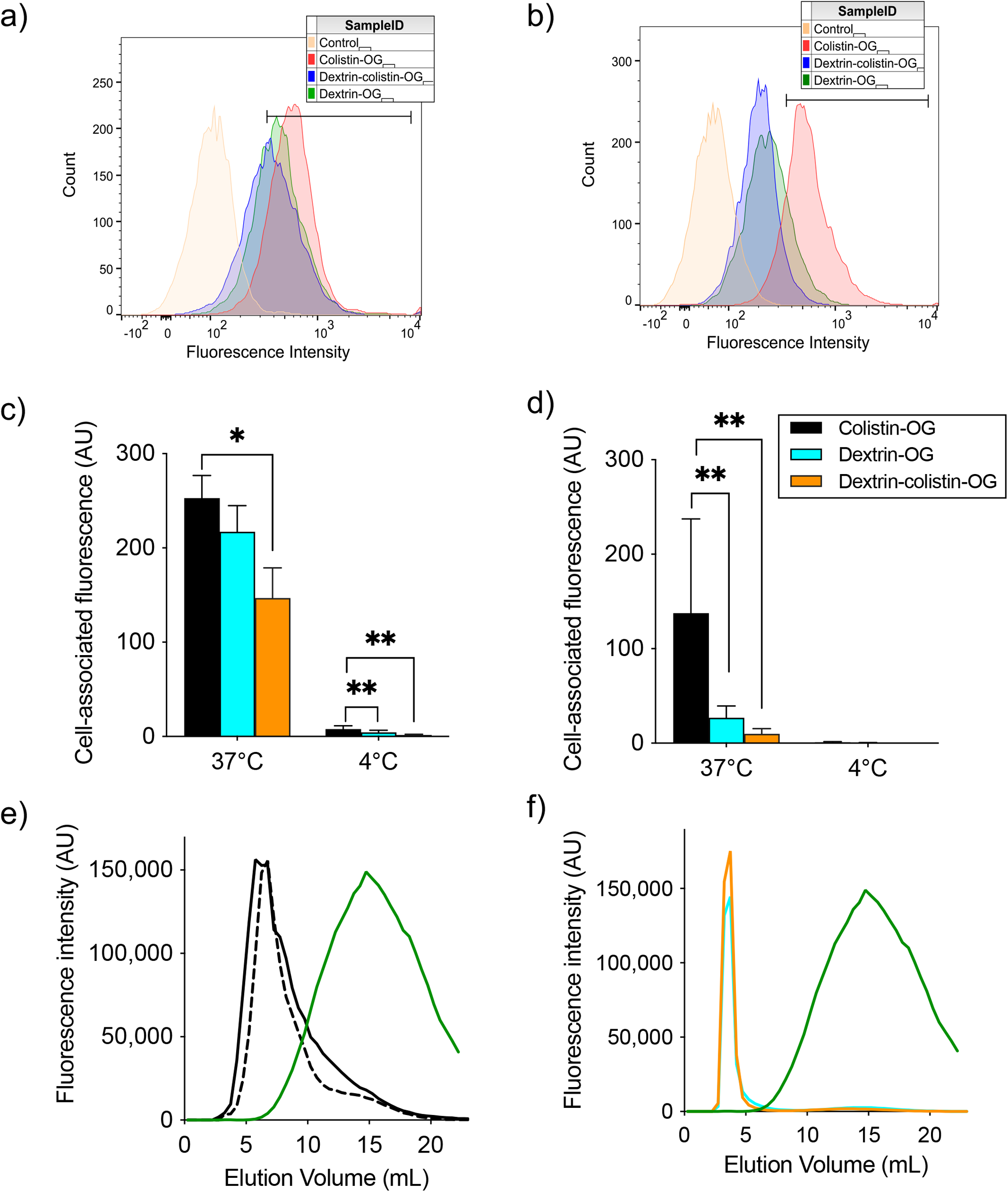
Binding and internalisation of OG cadaverine (control) and OG-labelled colistin, succinoylated dextrin and dextrin-colistin conjugate by (a,c) HK-2 and (b,d) NRK-52E cells after 1 h incubation at 4 and 37°C. Panels a) and b) show representative histograms for cell-associated fluorescence at 37°C. Panels c) and d) show cell-associated fluorescence at 37°C (total association) and 4°C (external binding) of OG-labelled colistin, dextrin and dextrin-colistin conjugates. Data are expressed as mean ± SEM (*n*=9). Significance vs. colistin-OG, where * = *p*<0.05 and ** = *p*<0.001. Panels e) and f) show PD-10 characterisation of cell culture medium after 4 h incubation of HK-2 cells with e) colistin-OG (solid line = pre-incubation, dashed line = post-incubation) or f) dextrin-OG (blue) and dextrin-colistin-OG (orange). OG cadaverine is shown by a green line. *OG* Oregon green, *AU* Arbitrary units, *C* Celcius, *mL* millilitre

**Figure 3.**
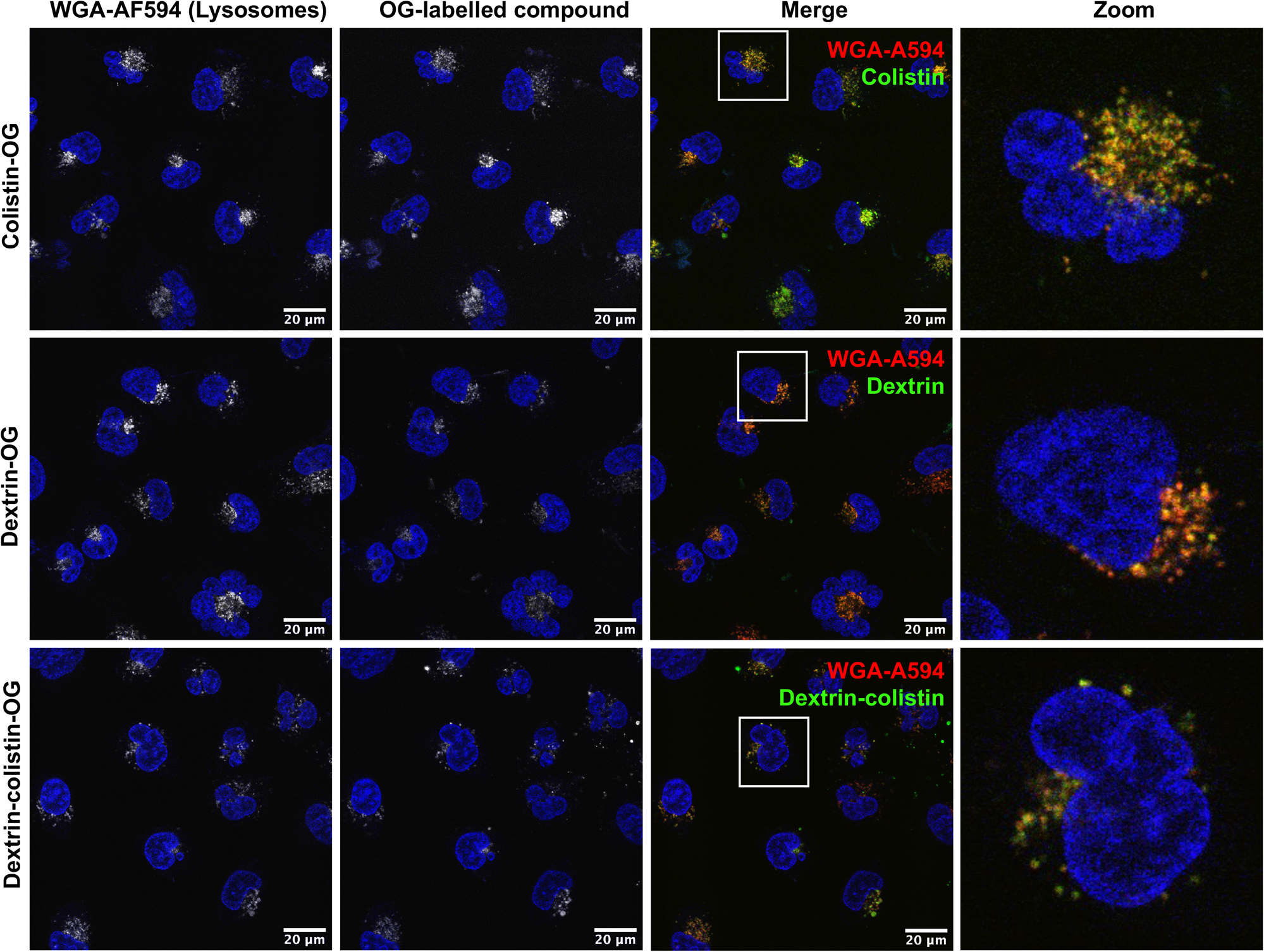
Uptake of OG-labelled colistin, dextrin and dextrin-colistin conjugate following a 2 h pulse and 16 h chase in HK-2 cells. WGA-AF594 was used to identify late endolysosomes and Hoechst 33342 (blue) was used as a nuclear marker. Scale bars show 20 µm. *WGA-A594* Wheat germ agglutinin-AlexaFluor 594,*OG* Oregon green

### *In vivo* acute kidney injury model

Mice treated with colistin sulfate exhibited weight loss (up to 6% of body weight) during treatment (Figure S8. In contrast, weight loss was absent in mice treated with dextrin-colistin conjugates; indeed mice treated with DC7 showed up to 3% weight gain. A statistically significant decrease in accumulation of dextrin-colistin conjugate in the kidney, liver and brain was observed (Figure 4), when compared to free drug (*p*<0.0001, 0.0001 and 0.01, respectively). Accumulation of both, colistin sulfate and dextrin-colistin conjugates was most pronounced in the kidney. Treatment with colistin sulfate induced significantly elevated levels of AKI biomarkers (KIM-1, NGAL) and urea (*p*<0.05), while no increases were observed in mice treated with dextrin-colistin conjugates (Figure 5a,b,c). None of the treatments significantly altered creatinine levels (Figure 5d). The kidneys of control mice treated with saline exhibited normal histology (Figure 6a, Table 3), whereas the kidneys of mice treated with colistin sulfate showed areas of mild tubular and interstitial damage (Figure 6b,c) including focal tubulointerstitial inflammation, haemorrhage (in <25% of tissue) and necrosis (Table 3). Interestingly, whilst minimal pathological changes were observed in the renal biopsies of mice treated with DC1 and DC2, mice treated with DC7 showed signs of thickening of the Bowman capsule and retraction of the glomerular tuft, indicative of mild glomerulonephritis (Figure 6d, Table 3).

**Figure 4.**
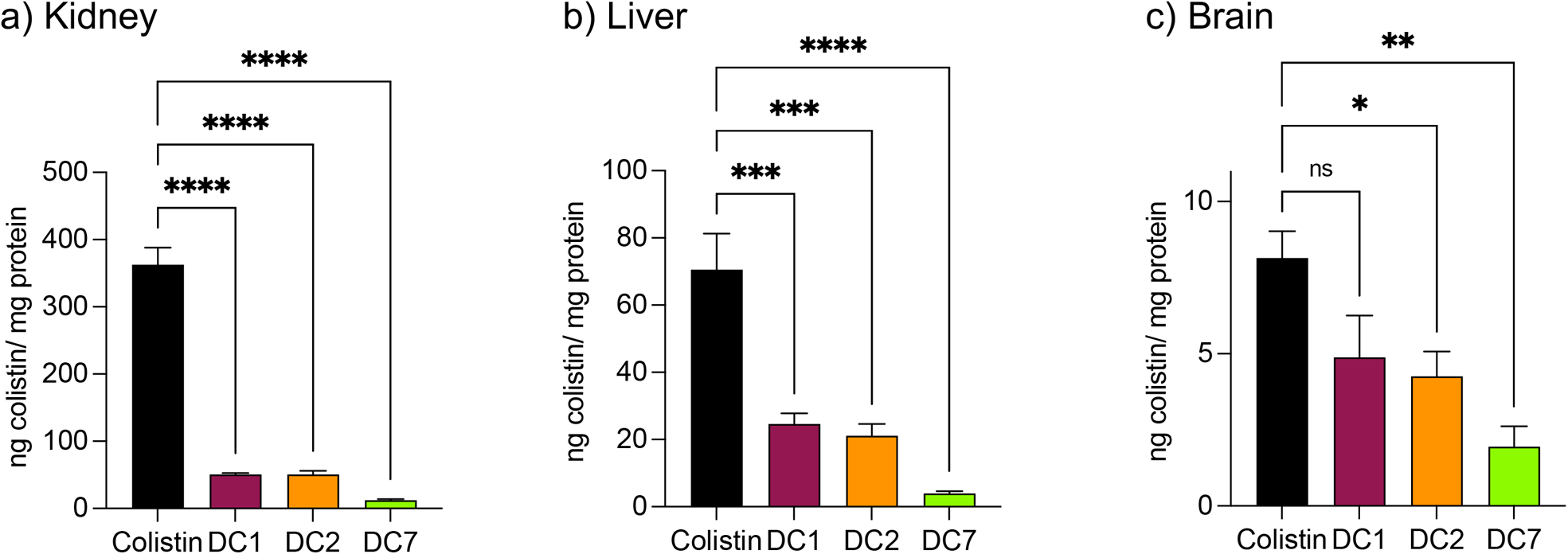
Accumulation of colistin in a) kidney, b) liver and c) brain in mice after twice-daily dosing for 7 days. Data are expressed as mean ±SEM (*n*=4-8). Significance vs. colistin-OG, where * = *p*<0.05, ** = *p*<0.01, *** = *p*<0.001, **** = *p*<0.0001 and ns = not significant. *ng* Nanogram *mg* Milligram

**Figure 5.**
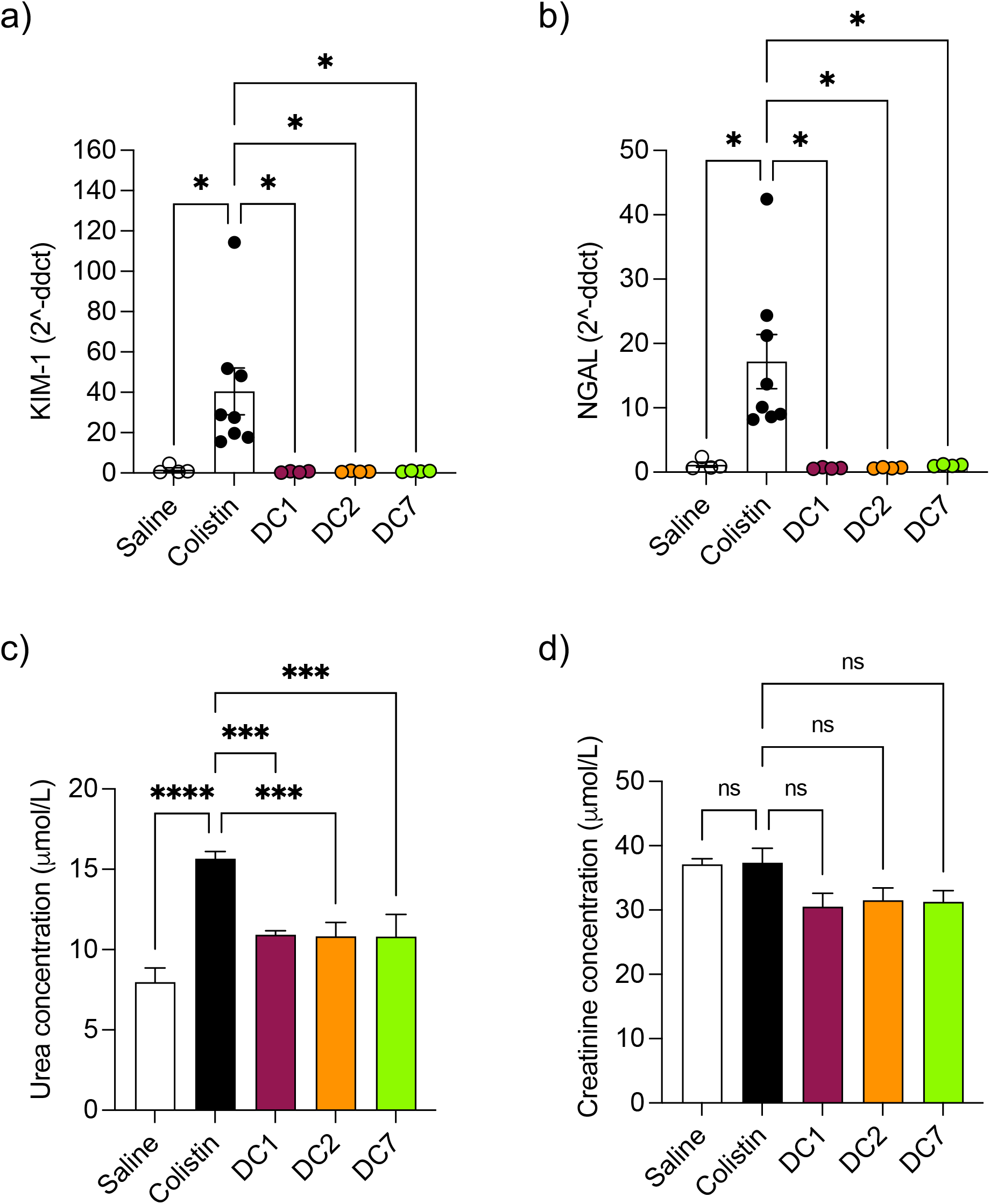
Detection of AKI biomarkers a) KIM-1, b) NGAL, c) urea and d) creatinine in mice after twice-daily dosing for 7 days. Data are expressed as mean ±SEM (*n*=4-8). Significance vs. colistin-OG, where * = *p*<0.05, *** = *p*<0.001, **** = *p*<0.0001 and ns = not significant. KIM-1 Kidney injury molecule-1, NGAL Neutrophil gelatinase-associated lipocalin, *ddct* Delta delta cycle threshold, *μmol/L M*icromole per litre

**Figure 6.**
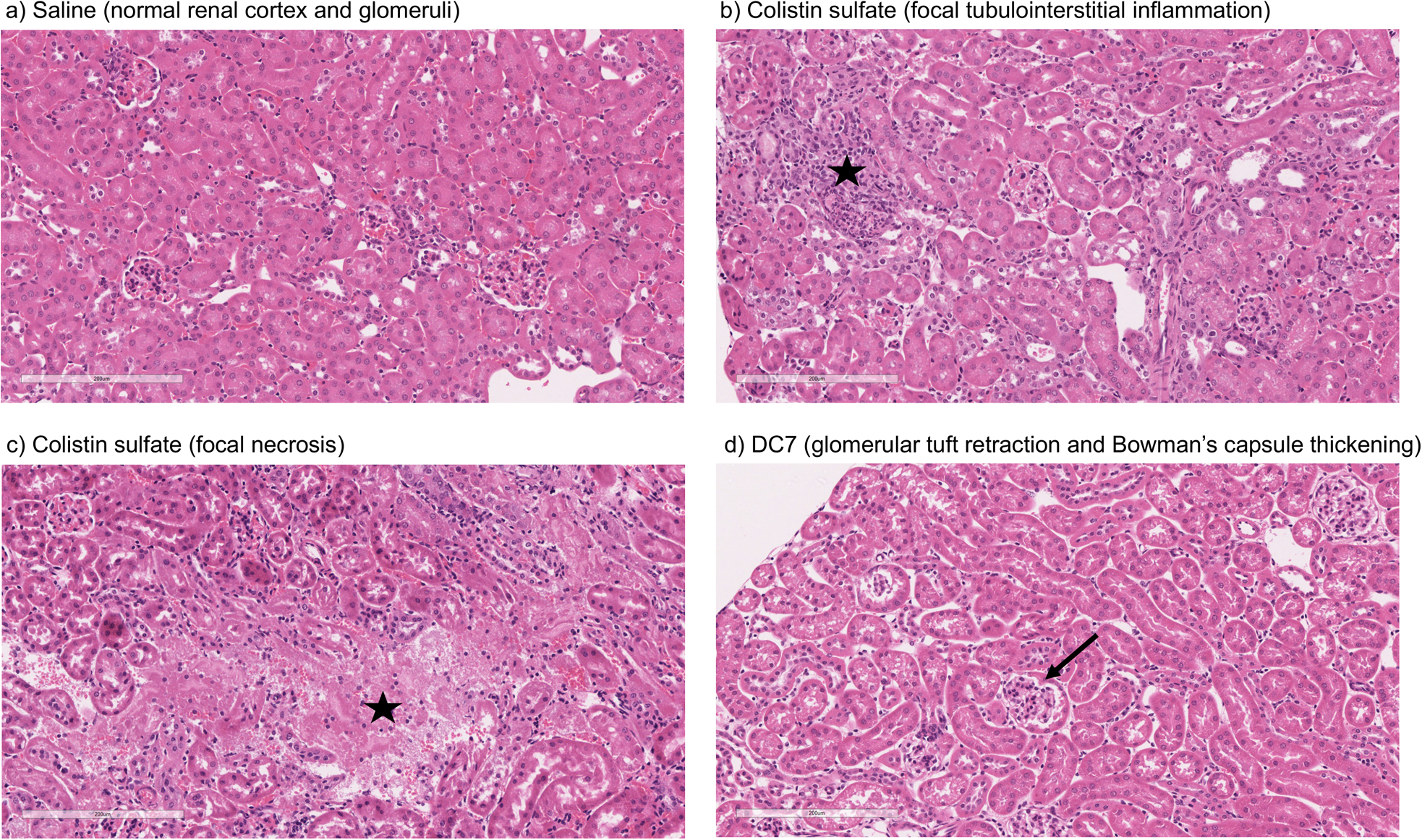
Representative images of kidney of mouse administered a) saline, b) colistin sulfate, c) colistin sulfate or d) DC7 for 7 days (scale bar = 200 μm), where ★ indicates areas of inflammation and necrosis, and ➝ indicates thickening of Bowman’s capsule.

**Table 3.** Semi-quantitative analysis of histological changes observed in renal biopsies of mice treated with saline (control), colistin sulfate or dextrin-colistin (DC) conjugates for 7 days (*n = 4-8*).

|  | Abnormality grade given to individual mice in each group |  |  |  |  |
| --- | --- | --- | --- | --- | --- |
|  | Control | Colistin sulfate | DC1 | DC2 | DC7 |
| Tubular | 0,0,0,0 | 0,0,0,0,0,0,0,0 | 0,0,0,0 | 0,0,0,0 | 0,0,0,0 |
| Endothelial | 0,0,0,0 | 0,0,0,0,0,0,0,0 | 0,0,0,0 | 0,0,0,0 | 0,0,0,0 |
| Glomerular | 0,0,0,0 | 0,0,0,0,0,0,0,0 | 0,0,0,0 | 0,0,0,1 | 1,0,2,2 |
| Tubulo/Interstitial | 0,1,1,0 | 0,0,1,1,0,0,0,2 | 0,0,0,0 | 0,0,0,1 | 1,0,0,1 |

Note: Histological damage graded according to the EGTI histology scoring system.<sup>21</sup>

## Discussion

Whilst acute kidney injury (AKI) remains a treatment-limiting adverse effect of colistin, its use has, however, resurged due to the emergence and spread of virulent multidrug-resistant (MDR) and extensively drug-resistant (XDR) Gram-negative bacilli, especially the ESKAPE pathogens, *Klebsiella pneumoniae, Acinetobacter baumannii, Pseudomonas aeruginosa*, and *Enterobacter* species.^22^

### Approaches to reducing polymyxin-induced AKI

Several approaches to preventing colistin’s dose-limiting nephrotoxicity have been investigated, including actions to promote the safe use of the antibiotic (e.g. identification of risk factors, development of optimised dosing strategies, therapeutic drug monitoring and early AKI detection criteria) and the use of colistin in combination therapy with other antibiotics.^23–26^ Co-administration of antioxidant and/or antiapoptotic compounds, such as melatonin, vitamin C and E, lycopene and curcurmin have also shown some beneficial effects in reducing apoptotic markers and histopathological changes in kidney tissue, although human studies are limited.^27, 28^ In parallel, researchers have developed next-generation polymyxin derivatives that are less nephrotoxic than the parent drug.^29–33^ For example, polymyxin B nonapeptide was found to be >50-fold less cytotoxic than polymyxin B toward HK-2 cells,^34^ however it is devoid of antibiotic activity.^35, 36^ In a more recent study, Li et al^37^ described a series of polymyxin analogues with 2-Thr and 10-Thr modifications, many of which showed retained or enhanced (up to 2-8-fold) antibacterial activity and lower toxicity toward HK-2 cells *in vitro* (up to 2-fold). Here, dextrin-colistin conjugates were at least 3-fold less cytotoxic than colistin sulfate, with only a modest reduction in antimicrobial activity.^15^ A major benefit of dextrin conjugation over these alternative approaches is the ability to utilise the Enhanced Permeation and Retention (EPR) effect to specifically target inflamed and infected tissues,^38^ thereby increasing the local concentration of antibiotic in the infected tissue, while preventing uptake and accumulation in the kidneys. In a recent study, nanoparticle accumulation in wounds was 25-fold higher than in normal, and increased a further two-fold when wounds were infected.^39^

### *In vitro* toxicity and intracellular localisation

Although polymyxin-induced AKI is widely reported, its underlying molecular mechanism is not completely understood. Studies addressing the intracellular fate of colistin have previously shown that exposure causes mitochondrial dysfunction and ER stress.^40–42^ Yun et al^2^ also observed partial co-localisation of regioselectively labelled monodansylated polymyxin B probes (MIPS-9543 and MIPS-9544) within the mitochondria and ER of NRK-52E cells. In our study, co-localisation of OG-labelled colistin or dextrin-colistin conjugates within the ER was not examined, however, we did not observe any co-localisation of probes within BODIPY TR ceramide- or MitoTracker-labelled compartments (Golgi and mitochondria, respectively) in HK-2 cells. In our studies, as observed by Jarzina et al^43^, human proximal kidney cells were significantly more sensitive to colistin sulfate than the rat cell line, NRK-52E. Interestingly, the authors also observed increased lysosomal disruption and significantly lower accumulation of colistin in rat cells, compared to human cells, which is consistent with the reduced internalisation and co-localisation of colistin to the endolysosomes of NRK-52E cells, compared to HK-2 cells, observed in this study. They hypothesised that human proximal tubular cells may have increased expression of endocytic receptors for colistin and/or endocytic activity, compared to rat proximal tubular cells, though this has not yet been proven. Surprisingly, despite rat cells being less sensitive to colistin sulfate than human cells, dextrin-colistin conjugates behaved similarly in both cell lines. Analysis of the metabolic effects of dextrin and succinoylated dextrin revealed similar patterns of cytotoxicity in NRK-52E cells, but showed minimal toxicity in HK-2 cells (Figure S9), suggesting that the dextrin carrier may cause enhanced toxicity in rat cells. The correlation between intracellular polymyxin B concentration and toxicity has also been shown by Ahmed et al.^44^ They observed that polymyxin B caused more cell death and substantially more polymyxin B accumulation in HK-2, compared to A549 cells. The differences in colistin’s cytotoxicity observed here and in other studies indicate species- and/or cell type-specific differences in its activity. In our studies, caspase 3/7 activity after 24 or 72 h exposure generally reflected cell viability, rather than being indicative of a modified level of apoptosis activation; the induction of caspase-mediated apoptosis following colistin exposure has been observed in several previous studies.^40, 42, 45, 46^ Lee et al^47^ showed an early, dose-dependent increase in caspase 3/7 activity after incubation of HK-2 cells with colistin sulfate for 6 h, suggesting that the time points used in our studies may not have captured the changes.

### Colistin-induced acute kidney injury

*In vivo* pharmacokinetics and toxicity of colistin have been extensively studied in rodent models. The model employed here was adapted from a previous study and designed to study histopathological changes and alterations in AKI biomarkers, in the absence of significant toxicity to the animals.^48^ Based on previous studies,^48–50^ we initially administered 10 mg/kg/dose colistin sulfate intraperitoneally twice daily for 3 days. Whilst no difference in nephrotoxicity was seen between animals treated with colistin sulfate and saline, studies have shown that nephrotoxicity is associated with cumulative dose and treatment duration.^11, 51, 52^ Here, increasing the dose to 20 mg/kg/dose and extending the treatment duration to 7 days (cumulative dose to 280 mg/kg), resulted in significant increases in kidney injury biomarkers and histopathological damage.

In selecting biomarkers for evaluation of magnitude of AKI beyond functional (creatinine) and histological (semi-quantitative scoring) we were guided by the work of The Predictive Safety Testing Consortium.^53^ The ideal characteristics identified by this group are that a biomarker identifies kidney injury early, reflects the degree of toxicity, displays similar reliability across multiple species, localises the site of kidney injury, tracks progression of injury and recovery from damage, is well characterised with respect to limitations of its capacities, and is accessible in readily available body fluids or tissues. While no biomarker yet exactly matches this paradigm, the two most closely approximating and therefore most commonly employed current AKI biomarkers are KIM-1 and NGAL, and we therefore quantified both in this study. Here, we found that colistin sulfate caused a significant increase in both, KIM-1 and NGAL levels, without any significant effect on creatinine levels (as corroborated using a Crystal Chem Mouse creatinine assay kit, *unpublished*). Similarly, when Luo et al^54^ evaluated KIM-1 and NGAL as early indicators of gentamicin-induced nephrotoxicity in rats, the AKI biomarkers increased before serum creatinine levels changed. Serum creatinine is commonly used to estimate glomerular filtration rate, for the detection of nephrotoxicity, however, elevated levels tend to reflect advanced damage to the kidney and, thus, is not useful for early detection of colistin-induced AKI.^41^ In this study, none of the dextrin-colistin conjugates caused any effect on KIM-1 or NGAL levels, suggesting that dextrin conjugation can prevent colistin-induced AKI. The mechanism for the reduced toxicity is still unclear; live cell imaging showed localisation of both, colistin and dextrin-colistin conjugates in endolysosomes of HK-2 cells, and to a lesser extent in NRK-52E cells, while flow cytometry showed reduced uptake of dextrin-colistin conjugates by HK-2 and NRK-52E cells. Our previous studies have shown that, even after complete degradation of dextrin by amylase, colistin retains linker residues with differing lengths of glucose units attached, which modify antimicrobial activity.^12^ Thus, we hypothesise that these dextrin-based structural changes alter the internalisation and accumulation of colistin at the luminal side of renal proximal tubule cells, possibly by reducing binding to megalin receptors, thereby lowering the colistin concentration inside these cells. Indeed, analysis of the colistin content in the kidneys showed significantly reduced levels in mice treated with dextrin-colistin conjugates, compared with colistin sulfate controls.

Although the transport of dextrin-colistin conjugates across the peritoneal membrane could, theoretically, be limited, due to their relatively large size (>30 kDa) our previous work clearly demonstrates the rapid degradation of dextrin and reduction of molecular mass by physiological concentrations of amylase and the rapid amylase-triggered release of colistin from dextrin-colistin conjugates (and diffusion across dialysis membranes) in both, physicochemical and functional/ antimicrobial assays.^14, 15^ In support of this, increased bioavailability following IP administration of nanoformulated drugs, in comparison to the low molecular weight (parent) drug has been shown for PEG (20 kDa) IFN-beta-1a (being almost 100%)^55^ and also in Bac_E_-PEG (24 kDa)-Alexa conjugates, which have wider distribution than the unPEGylated peptides.^56^ Moreover, experimental studies indicate that IP administered plasma proteins (e.g. albumin (66.5 kDa), transferrin (79.5 kDa) and immunoglobulin G (150kDa)) and are readily absorbed across the peritoneal membrane and distributed like IV-administered plasma proteins.^57^

Here, mice treated with colistin sulfate showed signs of focal tubulointerstitial inflammation. A similar study in rats found no evidence of inflammation after twice daily treatment for 7 days,^8^ however they used colistimethate sodium, rather than colistin sulfate, which is a less toxic prodrug of colistin. Histological analysis of renal tissue from mice treated with DC7 showed signs of mild glomerulonephritis. We propose that this could be caused by accumulation of DC7 in these animals, resulting in cumulative osmotic effects. We have previously shown that the amylase-dependent degradation of DC7 is related to the degree of succinoylation, resulting in a prolonged plasma half-life of 21.2 h after a single intravenous dose.^15^ Thus, 12-hourly dosing would not allow complete clearance of DC7 between doses and would cause accumulation of the conjugate over the 7 days. In clinical practice, polymer conjugates are typically administered with dosing intervals of several days and even commercial liposomal antibiotic formulations to treat bacterial and fungal infections are usually administered daily. The dosing schedule for individual dextrin-colistin conjugate formulations will require optimisation in future experiments, but it is likely that they will each require a different dosing interval to optimise the dose-response relationship.

## Conclusions

This study shows that dextrin conjugation can effectively overcome the renal toxicity associated with the administration of colistin and shows the potential of using polymer conjugation to improve the side effect profile of nephrotoxic drugs. Our findings shed new light on colistin’s underlying molecular mechanism that warrants further investigation to optimise dextrin-colistin conjugates’ design and dosing schedule. Extensive *in vivo* work is now needed, both to decipher the mechanism of reduced AKI by dextrin conjugation and ensure efficacy in an infection model.

## Acknowledgments

We acknowledge the technical cell culture support provided by Dr Sarah Youde and Dr Maria Stack and thank Dr Catherine Naseriyan from Central Biotechnology Services, Cardiff University for her technical expertise in FACS. We thank Thomas L. Williams for LC-MS technical support and gratefully acknowledge histology/staining services provided by Sue Wozniak and Hayley Pincott from the Cellular Pathology Department at the University Hospital of Wales (Cardiff and Vale University Health Board). Serum creatine and urea quantification was performed by Sarah James from the Biochemistry and Immunology Department at the University Hospital of Wales (Cardiff and Vale University Health Board).

## Funding

This research was funded by UK Medical Research Council (MR/N023633/1). This research was also funded, in part, by the Wellcome Trust (204824/Z/16/Z). For the purpose of open access, the author has applied a CC BY public copyright licence to any Author Accepted Manuscript version arising from this submission.

## Author contributions

ELF and DWT conceived the project. ELF, MV, SR, EJS, LN, C-TL, DJF, PRT, ATJ and DWT designed the study and MV, SR, EJS, LN, EC and NB performed the experiments. CT-L and PRT developed and optimised the in vivo model. AM, AJ and DJF analysed histology samples and interpreted the data. ELF, DWT and SR acquired the funding. ELF took the lead in writing the manuscript. All authors provided critical feedback and helped shape the research, analysis and manuscript.

## Transparency declarations

None to declare.

## Supplementary data

Supplementary Methods, Supplementary Results, Scheme 1 to 2, Table S1 and Figures S1 to S9 are available as Supplementary data at JAC Online.

## Notes

### Competing Interest Statement

The authors have declared no competing interest.

