## Supplementary methods & data for "Dextrin conjugation to colistin inhibits its toxicity, cellular uptake and acute kidney injury *in vivo*"

### Conjugation of colistin to dextrin ameliorates nephrotoxicity: exploration of cellular uptake and intracellular fate.

### METHODS

#### Synthesis and characterisation of OG-labelled probes

To prepare OG-labelled dextrin, succinoylated dextrin (325 mg, 2.5 mol%) was dissolved under stirring in PBS buffer (3 mL, pH 7.4) in a 10 mL round-bottomed flask. To this, EDC (38.5 mg) and sulfo-NHS (43.6 mg) were added and the mixture stirred for 15 min before addition of OG cadaverine (10 mg, from a stock solution of 100 mg/mL in anhydrous DMF, stored at -20°C until use). The reaction mixture was left stirring in the dark for 5 h prior to purification by size exclusion chromatography (SEC, disposable PD-10 desalting column containing Sephadex G-25 (Cytiva, Little Chalfont, UK)).

To prepare the OG-labelled dextrin-colistin conjugate, a dextrin-OG intermediate was first prepared, as described above, and then conjugated to colistin and purified by FPLC, as described previously,[16, 17] The total OG content of OG-labelled dextrin and dextrin-colistin conjugates was determined spectrophotometrically by measuring absorbance at 485 nm. Free OG content was assessed by measuring fluorescence ($\lambda_{ex}$ = 485 nm, $\lambda_{em}$ = 520 nm, gain 1000) of fractions (1 mL) eluting from a PD-10 column, as described previously.[16, 17]

To prepare OG-labelled colistin, the antibiotic (13.8 mg) was dissolved under stirring in PBS (2.5 mL, pH 7.4) in a 10 mL round-bottomed flask. To this, OG carboxylic acid succinimidyl ester (5 mg, from a stock solution of 2 mg/mL in anhydrous DMF, stored at -20°C until use) was added dropwise. Then, the reaction mixture was left stirring in the dark at room temperature for 2 h. The solution was then purified by FPLC (ÄKTA, Amersham Pharmacia Biotech, UK) using a prepacked Superdex 30 26/600 column coupled with a UV detector set at 219, 280 and 470 nm. The reaction mixture was injected into a 5 mL loop and eluted using 0.1 M ammonium acetate (pH 6.9, 0.22 µm filter-sterilised) at a flow rate of 2.5 mL/min. To verify purity, fractions (15 mL) containing colistin-OG (typically between 230 and 290 mL) were analysed by HPLC-fluorescence, using a Dionex ICS-3000 ion chromatography system (Thermo Scientific, Gloucester, UK) equipped with a Dionex RF-2000 fluorescence detector set at $\lambda_{ex}$ = 500 nm, $\lambda_{em}$ = 524 nm, and a Dionex AS autosampler, to quantify free OG. Data was collected and processed using Chromeleon 6.8 software. Fractions were diluted 10x using 0.1 M ammonium acetate then separation was achieved using an XSelect CSH C18 column (130 Å, 3.5 µm, 3.0 × 150 mm) (Waters, Wilmslow, UK) connected to an XSelect guard column (130 Å, 3.5 µm, 2.1 × 5 mm) inside a column oven at 40°C. A binary linear gradient method was used (95 to 5% A over 15 min; flow rate 0.8 mL/min; injection volume 10 µL) where mobile phase A is water (0.1% of formic acid), and mobile phase B is acetonitrile (0.1% of formic acid). Finally, fractions of colistin-OG conjugate that did not contain free OG were pooled and desalted by repeated freeze-drying (x 5) to remove ammonium acetate. The final compound was stored at -20°C until use. Analysis of colistin-OG was additionally performed using LC-MS on a Synapt G2-Si quadrupole time-of-flight (QTOF) mass spectrometer (Waters, UK), operating in the positive electrospray ionization mode, coupled to an ACQUITY H-Class UPLC system (Waters, Wilmslow, UK). Separation was accomplished using an ACQUITY UPLC BEH column (1.7 µm, 2.1 x 100 mm, Waters) inside a column oven at 40°C. A multistep gradient method was used (0-2 min, 98% A; 2-20 min, 2% A; flow rate 0.3 mL/min), where mobile phase A is water (0.1% formic acid), and mobile phase B is acetonitrile (0.1% formic acid).

### METHODS

#### Evaluation of *in vitro* cytotoxicity

Cells were seeded into sterile black, clear-based 96-well microplates (HK-2 at 2,500 cells/well, NRK-52E at 3,000 cells/well) in 0.1 mL of complete media (CM; K-SFM with EGF and BPE or DMEM with 5% v/v FBS, respectively). Cells were allowed to adhere for 24 h at 37°C with 5% $CO_2$ for 24 h. Filter-sterilised (0.22 µm) stock solutions of test compounds were prepared in PBS and used to supplement CM, which was used to replace the medium in the wells of the microplate. Test compounds were evaluated in triplicate at concentrations up to 1 mg/mL colistin base. Vehicle-only (PBS), apoptosis (1-10µM staurosporine) and necrosis (100µM Triton-X100) controls were also included. After a further 24 or 72 h incubation at 37°C with 5% $CO_2$, plates were processed in the dark as follows.

To measure necrosis, 25 µL of supernatant from each well was transferred to a fresh black 96-well microplate, containing 25 µL of CytoTox-One™ assay reagent. Plates were gently mixed then incubated at 21°C for 10 min, then a "stop solution" was added and fluorescence measured immediately at $\lambda_{ex}$ = 560 nm, $\lambda_{em}$ = 590 nm using a Fluostar Omega microplate reader. Total cellular LDH activity was measured after the addition of LDH lysis solution to cells, prior to addition of the assay reagent. Next, to measure cytotoxicity, CTB reagent was added to the wells of the original microplate ((10 µL/well). Plates were gently mixed then incubated at 37°C with 5% $CO_2$ for 1 h and fluorescence measured at $\lambda_{ex}$ = 560 nm, $\lambda_{em}$ = 590 nm using a Fluostar Omega microplate reader. Finally, 60 µL of Caspase-Glo 3/7 assay reagent was added to each well of the original microplate. Plates were gently agitated then incubated at room temperature (20°C) for 1 h, then luminescence measured using a Fluostar Omega microplate reader. Each cell line was plated in triplicate (n=3 technical replicates) and each experiment was repeated twice (n=2 biological replicates). Data was corrected for no-cell background, then expressed as percentage of the response of vehicle-only control cells (mean ± SD).

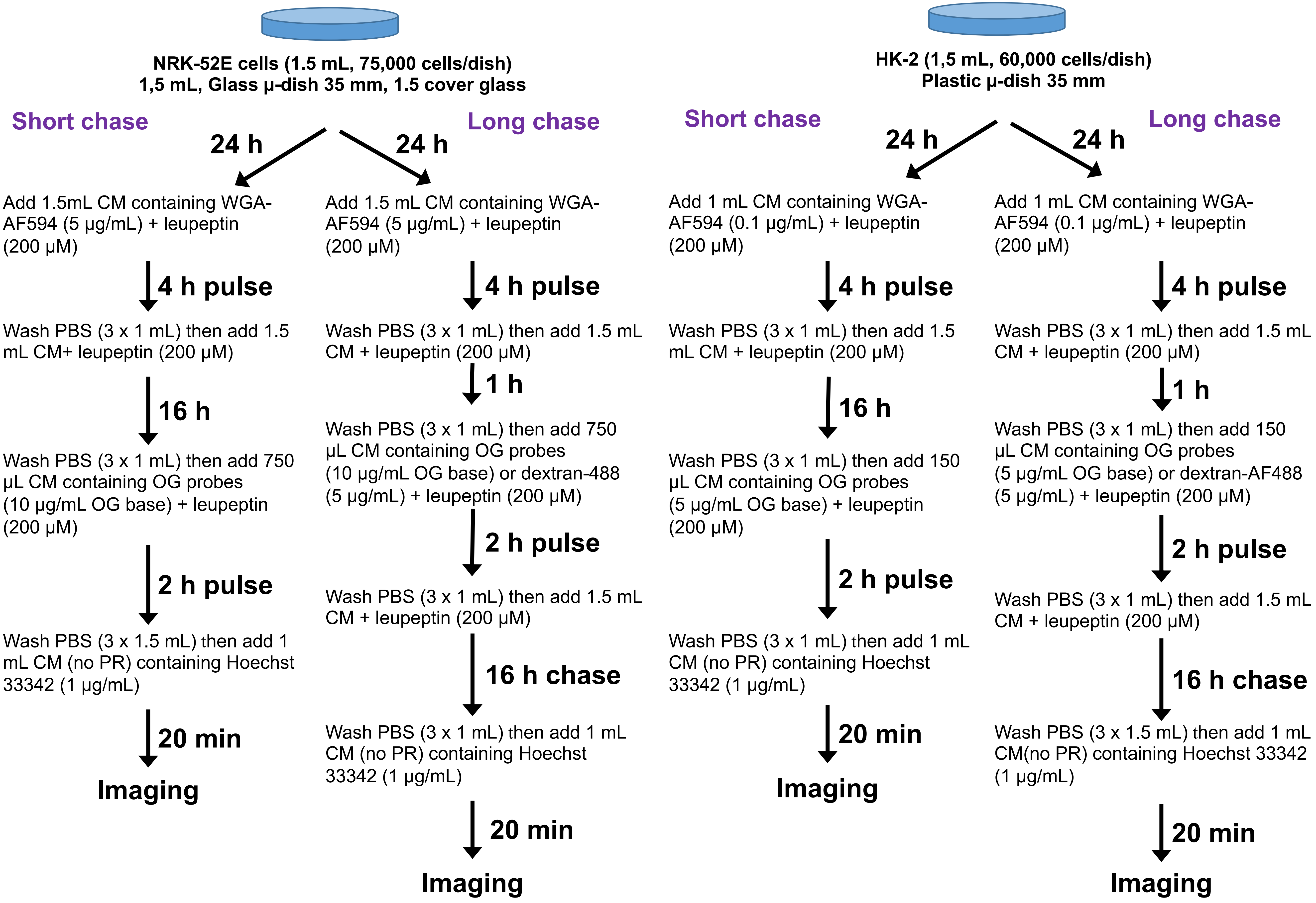


**Scheme S1.** Schematic describing the protocols used for confocal microscopy experiments for studying localisation of dextran-AF488 (control) and OG probes in endolysosomes in HK-2 and NRK-52E cell lines. (CM = complete medium, PR = phenol red)

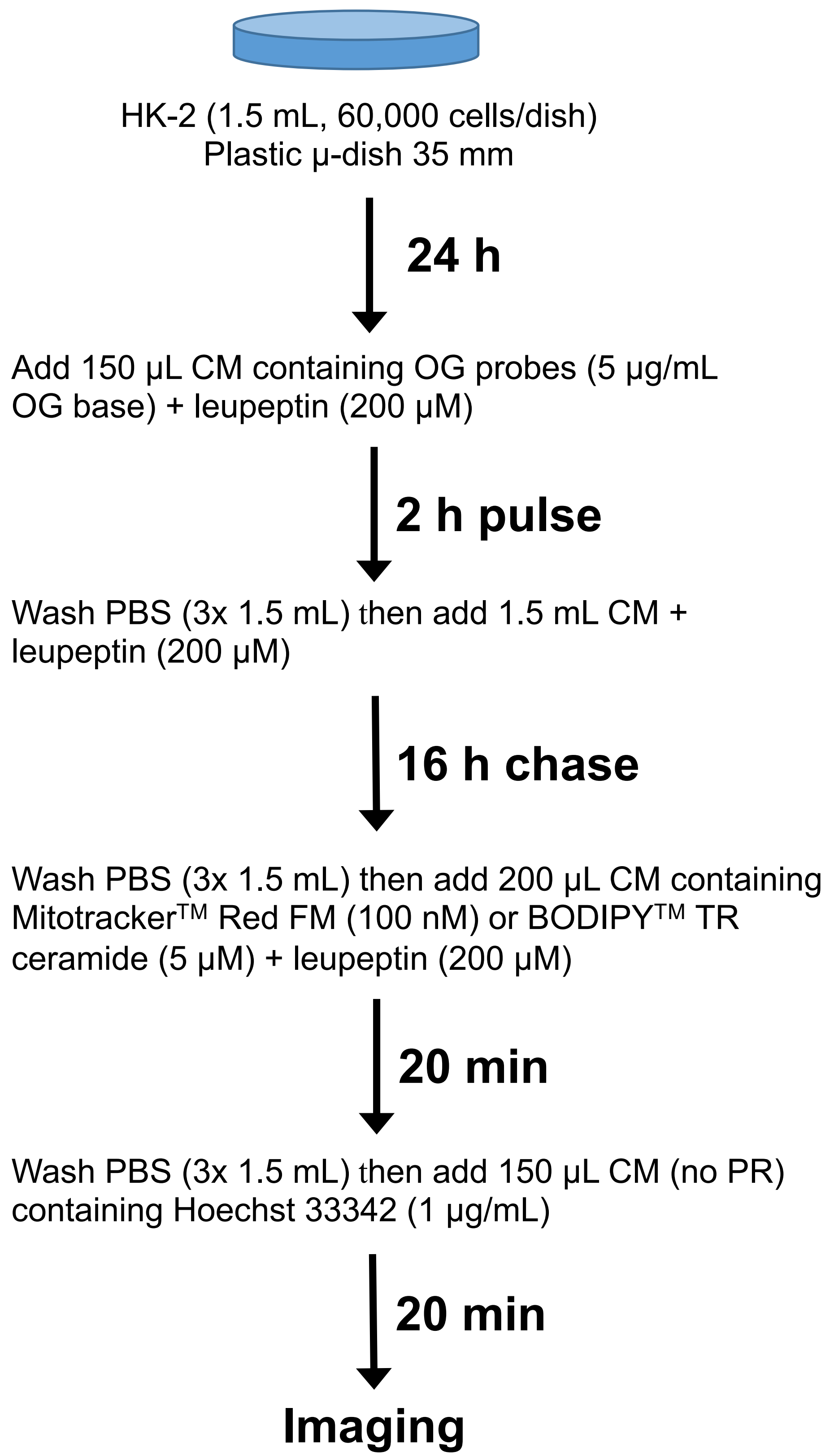


**Scheme S2.** Schematic describing the protocols used for confocal microscopy experiments when studying localisation of OG probes in mitochondria and Golgi in HK-2 cells. (CM = complete medium, PR = phenol red)

**Table S1.** Sequences of primers used for RT-qPCR.

| Gene | Sequence | Product (bp) |
|---|---|---|
| **KIM-1** | ACCGTGGCTATCACCAGGTA<br>TTCAGCTCGGGAATGCACAA | 123 |
| | CTGCTGCTACTGCTCCTTGT<br>GGAAGGCAACCACGCTTAGA | 91 |
| **NGAL** | GCTGTCGCTACTGGATCAGA<br>CTGTACCTGAGGATACCTGTGC | 89 |
| | GGAGCGATCAGTTCCGGG<br>CTGATCCAGTAGCGACAGCC | 181 |

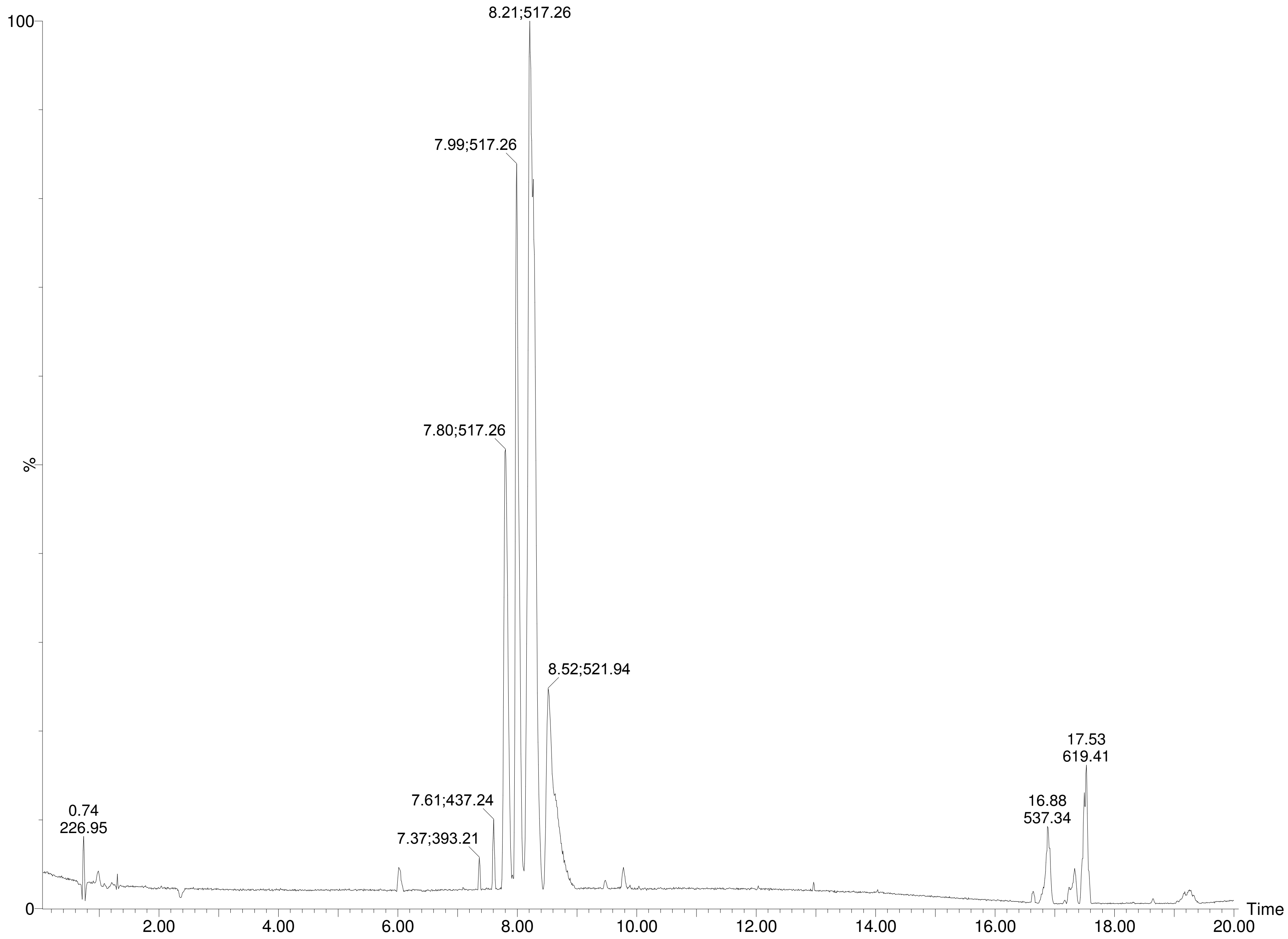


**Figure S1.** UPLC-QTOF-MS chromatograms (Base Peak Intensity, BPI) of colistin labelled with-Oregon Green (colistin-OG).

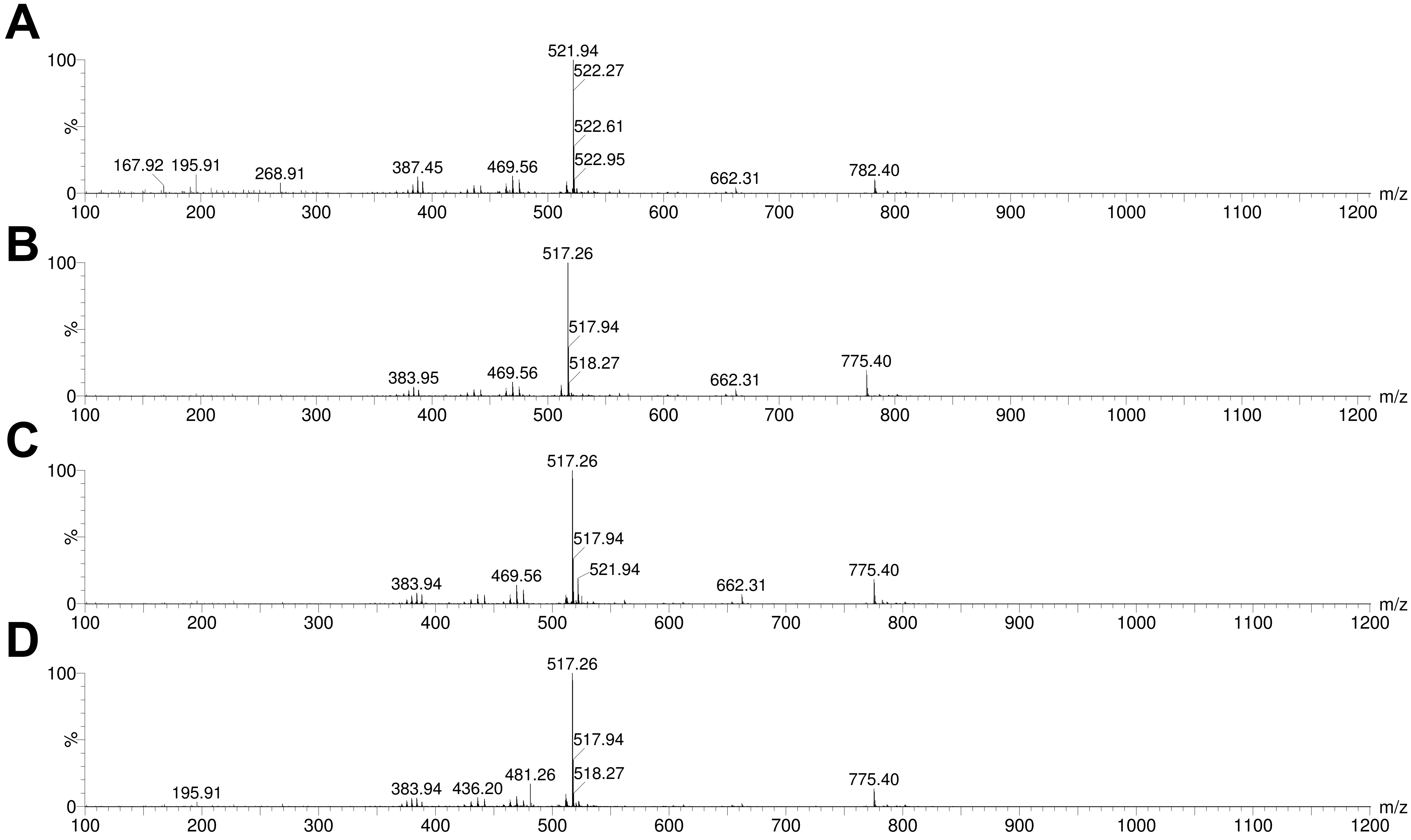


**Figure S2.** Q-TOF-MS spectra of the 4 main eluted peaks observed for colistin labelled with Oregon Green with a retention of A. 8.52 min, B. 8.21 min, C. 7.99 min and D. 7.80 min.

**Table S2.** Identification of the detected species from UPLC-QTOF-MS analysis of colistin labelled with Oregon Green.

| Retention time (min) | Observed mass | Calculated mass | Species | Adducts | Molecular formula |
|---|---|---|---|---|---|
| 7.80 | 517.26 | 517.27 | Colistin B-OG | $[M+3H]^{3+}$ | $C_{73}H_{106}F_2N_{16}O_{19}$ |
| 7.99 | 517.26 | 517.27 | Colistin B-OG | $[M+3H]^{3+}$ | $C_{73}H_{106}F_2N_{16}O_{19}$ |
| 8.21 | 517.26 | 517.27 | Colistin B-OG | $[M+3H]^{3+}$ | $C_{73}H_{106}F_2N_{16}O_{19}$ |
| 8.52 | 521.94 | 521.94 | Colistin A-OG | $[M+3H]^{3+}$ | $C_{74}H_{108}F_2N_{16}O_{19}$ |

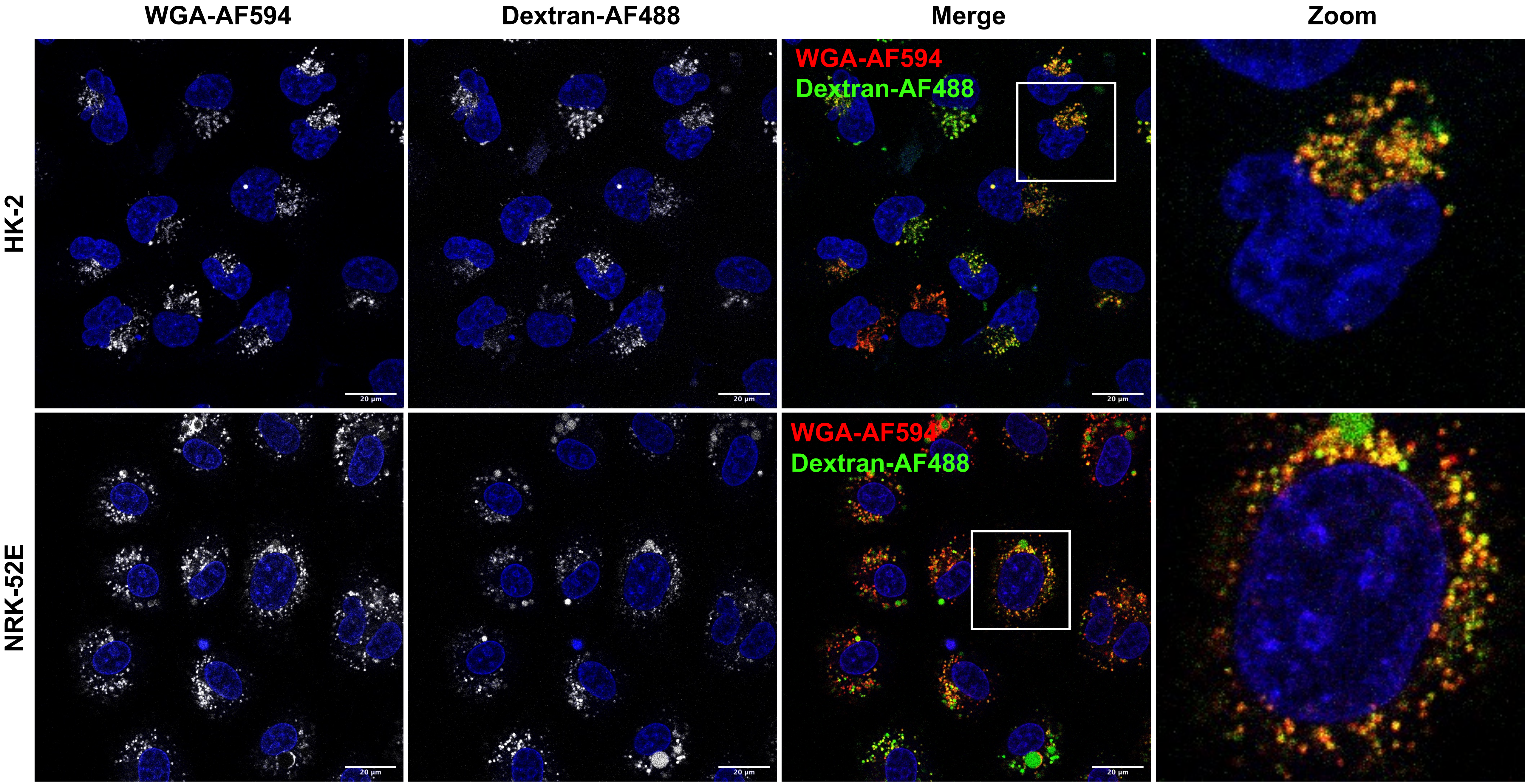


**Figure S3.** Uptake and distribution of WGA-AF594 and dextran-AF488 following a 4 and 2 h pulse, respectively, followed by a 16 h chase in HK-2 and NRK-52E cells. Hoechst 33342 (blue) was used as a nuclear marker. Scale bars show 20 µm.

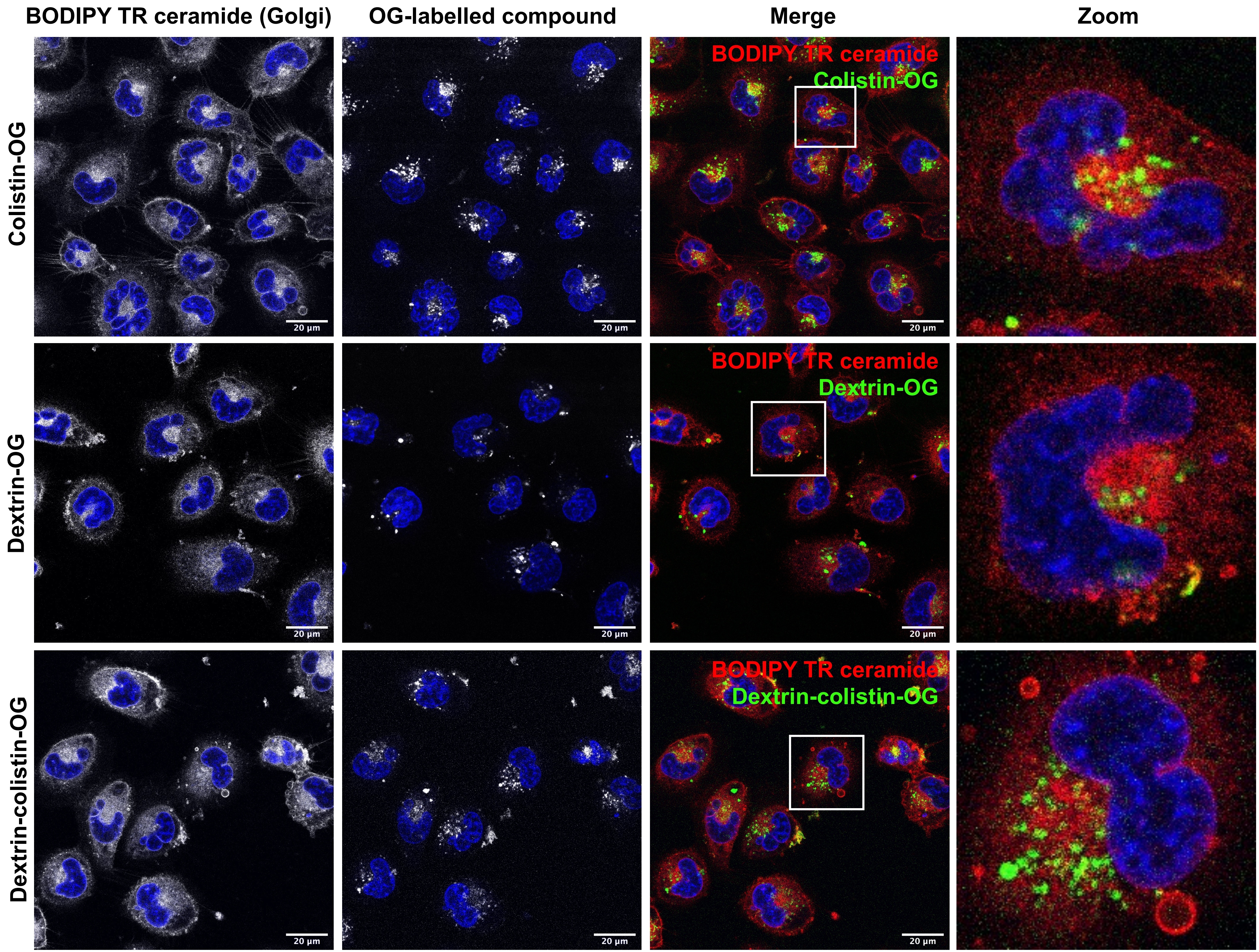


**Figure S4.** Uptake of OG-labelled colistin, dextrin and dextrin-colistin conjugate following a 2 h pulse and 16 h chase in HK-2 cells. BODIPY$^{TM}$ TR ceramide complexed to BSA was used to identify Golgi apparatus and Hoechst 33342 (blue) was used as a nuclear marker. Scale bars show 20 µm.

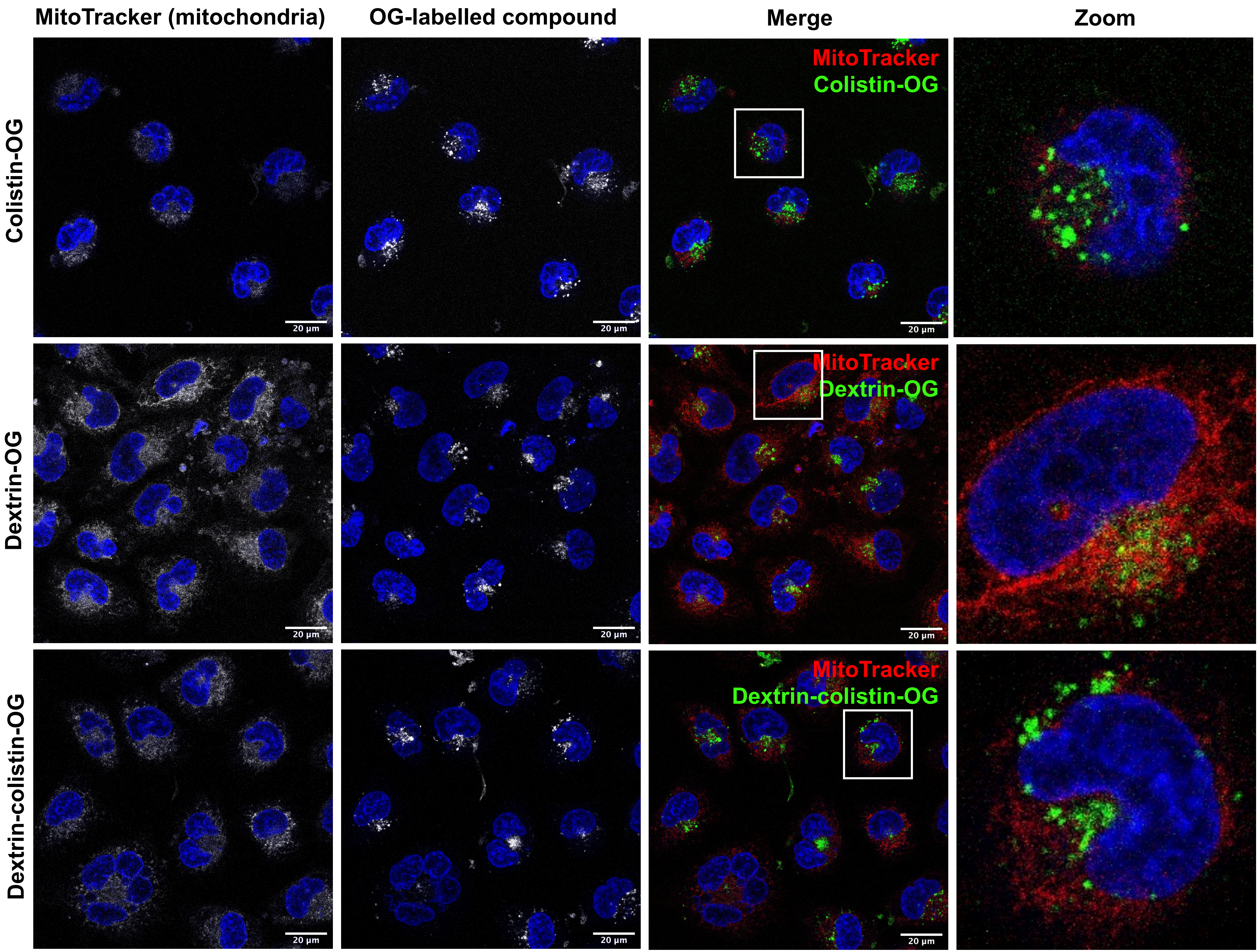


**Figure S5.** Uptake of OG-labelled colistin, dextrin and dextrin-colistin conjugate following a 2 h pulse and 16 h chase in HK-2 cells. MitoTracker Red FM was used to identify the mitochondria and Hoechst 33342 (blue) was used as a nuclear marker. Scale bars show 20 µm.

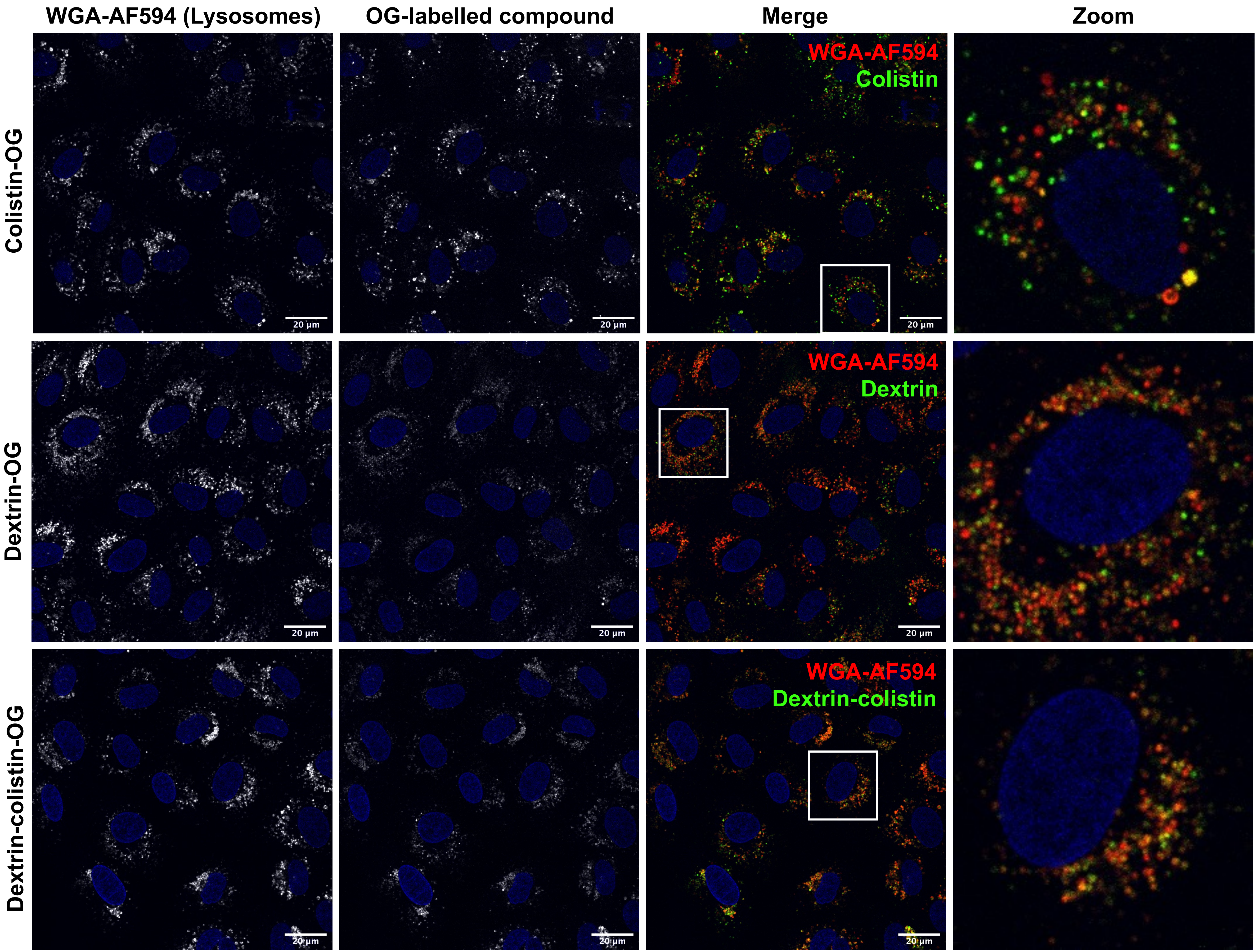


**Figure S6.** Uptake of OG-labelled colistin, dextrin and dextrin-colistin conjugate following a 2 h pulse and 16 h chase in NRK-52E cells. WGA-AF594 was used to identify late endolysosomes and Hoechst 33342 (blue) was used as a nuclear marker. Scale bars show 20 µm.

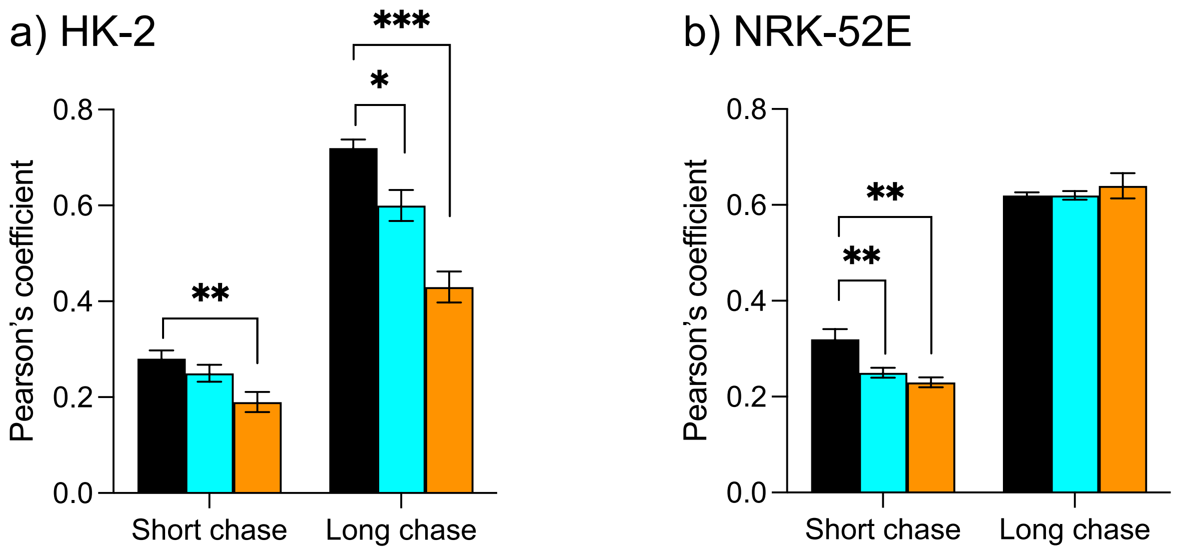


**Figure S7.** Pearson's coefficient characterising co-localisation of OG-labelled colistin (black), succinoylated dextrin (blue) and dextrin-colistin conjugates (orange) with WGA-AF594 after short and long chase in endo-lysosomal compartments of a) HK-2 and b) NRK-52E cells. Significance vs. colistin-OG, where * = $p$<0.05, ** = $p$<0.01, *** = $p$<0.0001.

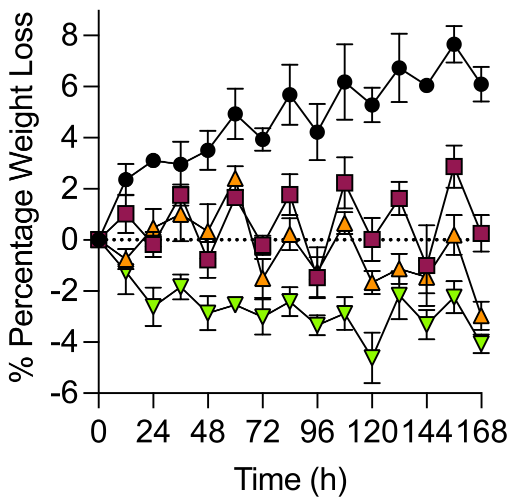


**Figure S8.** Percentage of body weight loss during treatment for 7 days with colistin sulfate (black), DC1 (red), DC2 (orange) and DC7 (green). Body weight was observed every 12 h.

**Method:**

Cells were seeded into sterile 96-well microplates (HK-2 at 2,500 cells/well, NRK-52E at 3,000 cells/well) in 0.1 mL of complete media and allowed to adhere for 24 h. The medium was then removed, and filter-sterilised test compounds were added to the wells. After a further 67 h incubation, MTT (20 µL of a 5 mg/mL solution in PBS) was added to each well and incubated for a further 5 h. The medium was then removed, and the precipitated formazan crystals solubilized by the addition of optical grade DMSO (100 µL). After 30 min, the absorbance of each well was measured at 540 nm using a microtiter plate reader. Cell viability was expressed as a percentage of the viability of untreated control cells and expressed as mean ± standard error of the mean (SEM) (*n* = 18).

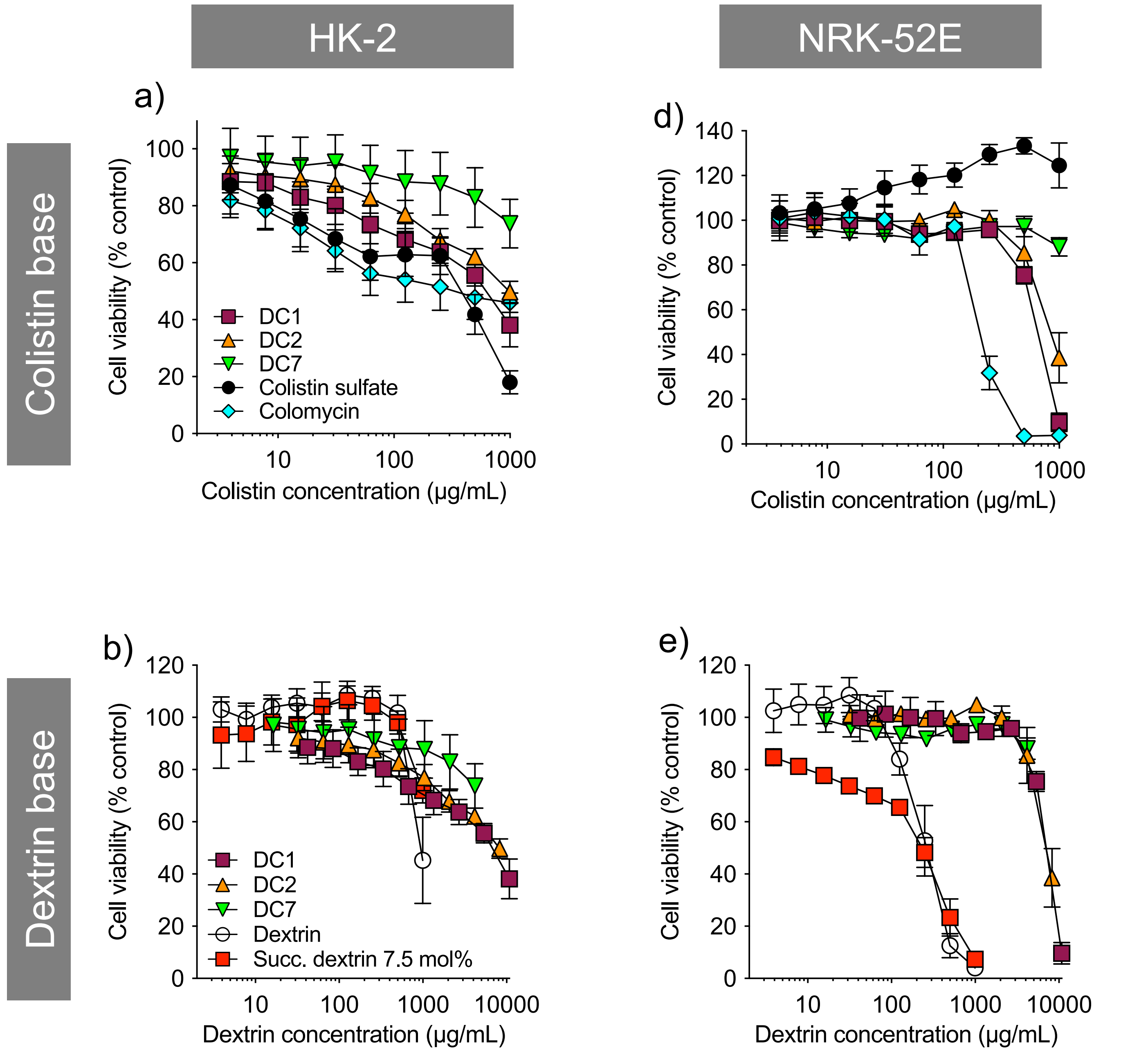


**Figure S9.** Metabolic activity (MTT assay) of (a,b) HK-2 and (c,d) NRK-52E cells incubated for 72 h with colistin sulfate, Colistimethate sodium (Colomycin®), dextrin-colistin conjugates, dextrin and succinoylated dextrin, as a percentage of untreated (media only) controls. Data represent mean ± SD (*n*=3). Where error bars are invisible, they are within the size of data points.
